# Different force fields give rise to different amyloid aggregation pathways in molecular dynamics simulations

**DOI:** 10.1101/2020.09.09.290320

**Authors:** Suman Samantray, Feng Yin, Batuhan Kav, Birgit Strodel

## Abstract

The progress towards understanding the molecular basis of Alzheimers’s disease is strongly connected to elucidating the early aggregation events of the amyloid-*β* (A*β*) peptide. Molecular dynamics (MD) simulations provide a viable technique to study the aggregation of A*β* into oligomers with high spatial and temporal resolution. However, the results of an MD simulation can only be as good as the underlying force field. A recent study by our group showed that none of the force fields tested can distinguish between aggregation-prone and non-aggregating peptide sequences, producing the same and in most cases too fast aggregation kinetics for all peptides. Since then, new force fields specially designed for intrinsically disordered proteins such as A*β* were developed. Here, we assess the applicability of these new force fields to studying peptide aggregation using the A*β*_16*−*22_ peptide and mutations of it as test case. We investigate their performance in modeling the monomeric state, the aggregation into oligomers, and the stability of the aggregation end product, i.e., the fibrillar state. A main finding is that changing the force field has a stronger effect on the simulated aggregation pathway than changing the peptide sequence. Also the new force fields are not able to reproduce the experimental aggregation propensity order of the peptides. Dissecting the various energy contributions shows that AMBER99SB-*disp* overestimates the interactions between the peptides and water, thereby inhibiting peptide aggregation. More promising results are obtained with CHARMM36m and especially its version with increased protein–water interactions. It is thus recommended to use this force field for peptide aggregation simulations and base future reparameterizations on it.

## 1 Introduction

Intrinsically disordered proteins (IDPs) are a class of proteins which are either completely unstructured or contain large disordered regions in their native state, which comes with a high tendency towards self-assembly leading to non-toxic or toxic aggregates and fibrils that may be related to diseases.^1^ One of the IDPs is the amyloid-beta peptide (A*β*), forming the core of senile plaques inside the human brain, which is considered to be the hallmark of Alzheimer’s disease (AD).^2^ Decoding the conformational dynamics of IDPs is a critical step in understanding the process of protein aggregation and fibrillation.^3^ However, the structural flexibility and high aggregation propensity impede experimental investigations to capture the dynamics of IDPs at atomic level.^4^ An alternativ approach to experiments is provided by molecular simulations, which allow for the necessary temporal and spatial resolution to follow the motions of IDPs.^5,6^ The recent years have seen the development of force fields (FFs) that allow the reliable modeling of IDPs using molecular dynamics (MD) simulations.^6–10^ The flexible nature of IDPs necessitated the FFs originally developed for folded proteins to be revised in order to accurately characterize the unfolded protein state.

However, while numerous FFs have been developed and benchmarked for IDPs (see ref 11 and references therein), it remains to be shown that they can also capture their aggregation behavior correctly. Our group compared the performance of several FFs for the formation of hexamers of the A*β*_16*−*22_ peptide, which is the sequence ^16^KLVVFAE^22^ of A*β*, and mutants of this peptide.^12^ This benchmark included FFs from AMBER, CHARMM, GROMOS, and OPLS. One of the main conclusions was that GROMOS54A7^13^ and OPLSAA^14,15^ overstabilize protein–protein interactions, leading to an overestimation of the aggregation speed and an inhibition of protein-aggregate dissociation. Thereafter, Derreumaux and coworkers investigated the protein aggregation behavior for the A*β*_16*−*22_ dimer using 17 different FFs in combination with conventional MD simulations,^16^ following up their previous work where they had employed replica exchange MD to study dimers and trimers of A*β*_16*−*22_.^17^ They concluded that FFs from CHARMM with updated CMAP correction^18,19^ such as CHARMM22*,^20^ CHARMM36,^21^ and CHARMM36m,^9^ along with FFs based on AMBER99^22^ with modified torsional parameters for the backbone and for the four amino acids Ile, Leu, Asp, and Asn like AMBER99SB-ILDN^23^ and AMBER14SB^24^ are best suited for studying amyloid aggregation. From these FFs, Charmm36m is the only one developed for IDPs which was realized by refining backbone potentials in order to model the preference of IDPs to adopt extended structures.^9^ Recently, researchers at D. E. Shaw Research developed an FF aimed at being applicable to both folded and unfolded proteins leading to AMBER99SB-*disp*.^10^ To reach this goal, Robustelli at al. used AMBER99SB*-ILDNQ^23^ in combination with the TIP4P-D water model^8^ as starting point and introduced modifications to the backbone torsional potential and enhanced the interaction potential between the backbone carbonyl oxygens and backbone amide hydrogen atoms, which is thought to increase the overall stability of extended conformations as present in *β*-sheets.^10^ In addition, they revised the water model by increasing the C6 term determining the attractive part of the Lennard-Jones interactions, which is expected to avoid hydrophobic collapse of the proteins. An increased protein–water interaction usually leads to a stabilization of extended conformations.^8,25^

The tendency of A*β* towards aggregation has been proposed to result to a large extent from its hydrophobic core region (residues 17–21). While the A*β*_16*−*22_ segment is not sufficient to understand the aggregation of full-length A*β*, since the latter involves 5–6 times more residues compared to the former, which not only increases the conformational space but also influences the aggregation behavior, this short peptide is nonetheless an attractive model for studying amyloid aggregation. First, A*β*_16*−*22_ is able to form fibrils itself, which are characterized by an antiparallel ordering of the peptides forming the *β*-sheets.^26^ Second, given its small size it is ideally suited for exploring the thermodynamics and kinetics of its aggregation using experiments^27–29^ and also MD simulations. The rigorous mutation study by Senguen et al. showed that *π*–*π* interactions do not play an important role during A*β*_16*−*22_ aggregation.^27^ Instead, the hydrophobicity of the amino acids in the region 17–21 seems to be the dominant factor determining the aggregation speed of A*β*_16*−*22_ and mutants of it, while electrostatic contacts between K16 and E22 provide further stability of the A*β*_16*−*22_ aggregates and ensure proper orientation in antiparallel *β*-sheets.^30^ In the past two decades, numerous coarse-grained and all-atom MD simulations have been performed for studying the aggregation of this peptide into small oligomers. ^12,16,30–35^ For a comprehensive review of simulation studies of A*β*_16*−*22_ including both all-atom and coarse-grained peptide models, the reader is referred to ref 36.

However, a valid question is – considering the vastly different results that were obtained from FF benchmarks of A*β* ^37–40^ *and other IDPs*^*25*^ *– to what extent the simulation results of Aβ*_16*−*22_ aggregation are affected by the FFs used in these studies. This question is addressed in the current work, aiming to elucidate the FFs that are suited to study amyloid aggregation using the heptapeptide A*β*_16*−*22_ and two of its mutants as a test cases. Experiments by Senguen et al. showed that the single mutant F19L A*β*_16*−*22_ (*m*1) forms fibrils faster than wild-type A*β*_16*−*22_ (*wt*), while the double mutant F19V/F20V A*β*_16*−*22_ (*m*2) does no aggregate at all. ^27^ These findings can be explained with the increased and decreased hydrophobicity of *m*1 and *m*2, respectively. Since they were derived from sedimentation assays and dynamic light scattering that followed the monomer concentration of each of these peptide sequences until fibrils were formed,^27^ they allow us to compare our simulation results on small fibrillar oligomers with the experimental observations.^12^ In the first part of this study, we elucidate the structural transitions of the three peptide sequences at the monomeric level since it is known that the monomer state has an effect or even controls the aggregation process.^41–44^ Secondly, we investigate the formation of hexamers by these peptides and establish links between the monomer configuration and the oligomer state. To complement this view, we finally check the stability of a preformed steric zipper involving twelve copies of either *wt, m*1, or *m*2, providing a conclusive evidence regarding which of the FFs under study is best suited to study amyloid aggregation using MD simulations. The FFs included in our test set are AMBER99SB-*disp*, CHARMM36m and CHARMM36mW (which is based on CHARMM36m but includes more favorable van der Waals interactions between protein and water) recently developed for IDPs as well as the older force fields GROMOS54A7 and OPLS-AA, which are already known to overstabilize protein–protein interactions ^12^ but serve as a useful reference here.

## 2 Simulation details

### 2.1 Systems

To understand the influence of the different FFs on the kinetics and structures of amyloid aggregation, we simulated residues 16–22 of the amyloid-*β* peptide, considering the wild-type (*wt*) and the two mutant F19L (*m*1) and F19V/F20V (*m*2) sequences. We capped the N- and C-termini of the peptides with acetyl (ACE) and N-methlyamide (NME) groups, respectively, to mimic the experimental conditions.^27^ Each peptide was simulated using five different force fields with their respective water models: AMBER99SB-*disp* ^10^ with modified TIP4P-D^8^ (A99-d), CHARMM36m^9^ with TIP3P^45^ (C36m) and with increased protein– water interactions^9^ (C36mW), GROMOS54a7^13^ with SPC^46^ (G54a7), and OPLS-AA^14,15^ with TIP4P^47^ (OPLS). Throughout this paper, we will refer to each of these systems with their abbreviations given in the parentheses.

### 2.2 Monomer simulations

We simulated the monomeric peptides starting from extended states and solvated them with a cubic water box. We set the minimum distance between the peptides and the edges of the water box to 1.2 nm. Only for the translational diffusion simulations, we placed the monomers 1.7 nm away from the box edges, effectively doubling the simulation box volume. We assigned the ionization states of lysine and glutamic acid at pH 7 to be protonated and deprotonated, respectively, resulting in electrostatically neutral peptides. In all simulations, we added Na^+^ and Cl^−^ ions to reach an NaCl concentration of 150 mM. We minimized each system using the steepest descent algorithm, followed by equilibration, first with a 10 ps run in the *NV T* ensemble while constraining the heavy peptide atoms to their initial positions, afterwards with a 10 ps run in the *NpT* ensemble without position constraints. For the production runs we simulated each system for 500 ns in the *NpT* ensemble with *T* = 298 K and *p* = 1 bar. We obtained three independent trajectories for each system starting from the same initial structures but applying independent minimization and equilibration procedures. For the translational diffusion simulations we collected three 400 ns long trajectories for each system. Throughout all simulations we constrained all bond lengths using the LINCS algorithm.^48^ The electrostatic and van der Waals interactions were calculated using the particle mesh Ewald (PME) method^49^ and the real-space components truncated at 1.2 nm. We controlled the temperature and pressure using a velocity rescaling algorithm^50^ with a relaxation time of 0.1 ps and a Parrinello-Rahman barostat^51^ with a relaxation time of 2 ps, respectively. For the simulations with G54a7 and OPLS we employed virtual sites for the nonpolar hydrogen atoms^52^ allowing for an integration time step of 4 fs, while a time step of 2 fs was used for the remaining FFs.

### 2.3 Hexamer simulations

We introduced six peptides into a cubic box with 10 nm edge length to study the oligomer formation for each of the three peptides and FFs. The initial configurations for these simulations were generated with the software PACKMOL^53^ using the most populated peptide structures identified in the monomer simulations. We positioned the monomers in such a way that none of the monomer–monomer distances with respect to any atom pair was smaller than 0.4 nm or greater than 1 nm. The simulations were set up using the same procedure as described for the monomer systems in section 2.2. For A99-d, C36m, and C36mW we obtained three independent trajectories of 1 *µ*s length each, while for G54a7 and OPLS we used the existing data from previous simulations performed within our group,^12^ which contain five production runs in the *NpT* ensemble for 300 ns each.

### 2.4 Steric zipper simulations

In order to corroborate the results from the oligomer formation simulations, we tested the stability of preformed minifibrils composed of twelve peptides stacked in two layers with six peptides forming an antiparallel *β*-sheet in each layer (Fig. S1 in the Supporting Information). This arrangement is also called steric zipper.^54^ These minifibrils were generated using the microcrystal structure of KLVFFA as determined by X-ray crystallography (PDB code 3OW9)^55^ as starting point. After adding the E22 residue as well as the terminal capping groups ACE and NME to each of the twelve peptide chains, which was accomplished with PyMOL,^56^ the peptides had to be aligned again so that the terminal residues K16 and E22 were next to each other in the antiparallel *β*-sheet and above each other in the double layer. In the case of the mutants *m*1 and *m*2 the mutations F19L and F20V/F20V were introduced too. We placed each of the minifibrils in the center of a cubic water box of size 15 nm in each spatial dimension. We added Na^+^ and Cl^*−*^ to adjust the salt concentration at 150 mM. We performed the minimization, equilibration, and production runs with the same simulation parameters as described in section 2.2. For each system three production runs in the *NpT* ensemble and of 300 ns length were carried out. These simulations testing the stability of the steric zipper conformation were performed for A99-d, C36m, and C36mW.

### 2.5 Analysis

#### Structural characterization

The simulations were analyzed using a combination of standard GROMACS tools, VMD,^57^ and in-house Python scripts^58^ invoking the MDAnalysis^59^ and MDTraj^60^ libraries. We determined the representative monomer structures of the peptides using the Gromacs clustering tool of Daura et al.^61^ with a cutoff of the root mean square deviation (RMSD) of 0.2 nm. For the intra- and interpeptide contacts, we considered two residues to be in contact if the distance between any pair of atoms from residue *a* and residue *b* is 0.4 nm or less. Based on this distance cutoff also the size of the oligomers was determined. The nonbonded interaction energies consisting of van der Waals (vdW) and electrostatic interactions were calculated using the *rerun* option of GROMACS *mdrun* for all intra- and interpeptide residue–residue pairs. In order to calculate the interaction energies between peptides, we extracted the peptide pairs present in the simulations using the distance criterion of 0.4 nm or less between the two peptides. Thus, peptide pairs as present in dimers and higher-order oligomers up to hexamers are considered for this analysis. For the characterization of the steric zippers we calculated the number of hydrogen bonds (H-bonds) formed between the peptides within a *β*-sheet as well as the nematic order parameter, *S*_2_. An H-bond is defined to be formed if the donor–acceptor distance is less than 0.35 nm and the donor-H-acceptor angle is less than 30*°*. The nematic order parameter,

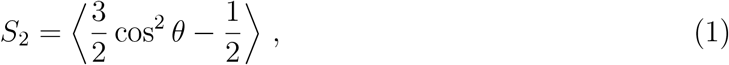

where the angle brackets refer to time and ensemble averaging and *θ* is the angle between the vector pointing from the N-terminus to the C-terminus of one peptide and the same kind of vector of another peptide in the system, was calculated with MDTraj. It describes the orientational order of a system with values ranging between 0 and 1 corresponding to isotropic and anisotropic systems, respectively.

#### Transition networks

In order to visualize the aggregation pathways, we constructed transition networks to elucidate the important intermediate stages.^12,34,62,63^ We define the states in the transition networks in terms of the oligomer size, which ranges from 1 for monomers to 6 for hexamers, and the *β*-sheet content divided into five ranges with 0–20%, 20–40% etc. In the resulting network models, which were plotted with Gephi,^64^ the nodes represent the intermediate aggregation states and the edges the transitions between these states. The size of the nodes are proportional to the population of the states and the edge thickness to the mass flux between the connected states. We calculated the *β*-sheet content based on the dihedral angles along the peptide backbone. A *β*-sheet is assumed to be formed when the *φ* and *ψ* values are located within the polygon with the vertices at (*−*180*°*, 180*°*), (*−*180*°*, 126*°*), (*−*162*°*, 126*°*), (*−*162*°*, 108*°*), (*−*144*°*, 108*°*), (*−*144*°*, 90*°*), (*−*50*°*, 90*°*), and (*−*50*°*, 180*°*).^30,34^ This definition allows to also assign a monomer to the *β*-state, which would not be possible if one used, as commonly done, the H-bond pattern between the peptides instead.

#### Translational diffusion

In order to accurately compute the translational diffusion constants of the A*β*_16*−*22_ peptide and its mutants, we extended the trajectories of the monomers to 400 ns in the *NpT* ensemble. It is important to note that the velocity-rescaling thermostat used in our simulations^50^ has been shown to produce transport properties indistinguishable from the *NV E* ensemble.^65,66^ Therefore, we did not run additional simulations in the *NV E* or *NV T* ensemble. We used the mean-squared displacement (MSD) and the Einstein relation in three dimensions to calculate the translational diffusion constants of the peptides,

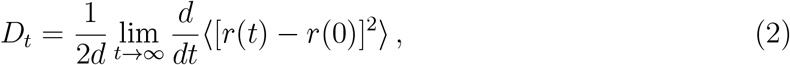

where *D*_*t*_ is the diffusion constant, *d* is the system dimension, *r*(0), *r*(*t*) are the particle positions at times *t* = 0 and *t*, and the term with angle brackets represents the MSD. For normal diffusion, the MSD grows linearly with sufficiently large values of time *t*. As a result, the three-dimensional translational diffusion constant *D*_*t*_ is equal to one sixth of the slope of the linear region of the MSD-vs-time curve. In order to obtain the MSD of the peptides, we divided each trajectory to 10 ns time windows with 5 ns lag time between two consecutive windows. This yields 80 subtrajectories per 400 ns trajectory, from which the MSD of the entire peptide was obtained by averaging over all atoms and subtrajectories. The three-dimensional translational diffusion constant *D*_*t*_ was calculated from the slope of the subtrajectory- and atom-averaged MSD curves by fitting a linear curve between 2 and 8 ns. The final translational diffusion constant *D*_*t*_ per system (i.e., per peptide and FF) and the associated standard error is reported as the average over three trajectories.

## 3 Results

### 3.1 Structural characterization of the A*β*_16*−*22_ monomer

When studying oligomer formation, it is important to separate the effects coming from the monomeric structure from those originating from interprotein interactions that drive the aggregation process. In this section, we assess how the different FFs affect the monomeric ensemble of A*β*_16*−*22_ based on RMSD-based clustering as well as intrapeptide contact maps and interaction energies.

#### Structural clustering and characterization

We performed an RMSD-based cluster analysis to explore the conformational heterogeneity in the ensemble of peptide structures generated by the MD simulations of a single *wt, m*1, and *m*2 peptide using the different FFs. By clustering structures that are within an RMSD of 0.2 nm, we calculated how the total number of observed clusters changes with time (Fig. 1). The plateau region of the number of clusters over time indicates that from *≈*300 ns on no new structures different from the clusters already identified were sampled, which applies to all FFs. The number of clusters obtained signify the unique structures sampled during the MD simulations. Irrespective of the peptide sequence the G54a7 force field yields the largest number of clusters, followed by OPLS, while C36m yields the lowest number of clusters. By counting the frequency of each cluster throughout the simulation trajectories, we observe that with A99-d, C36m, and C36mW the two dominant clusters are significantly more populated than the counterparts obtained with OPLS and G54a7. For instance, the first two clusters obtained with A99-d, C36m, and C36mW contain *>*85% of the structures for the *wt* peptide compared to 72% with OPLS and only 63% with G54a7 (the full list is given as Table S1 in the Supporting Information). This observation indicates that with the newer FFs the peptides are less flexible, which can result from increased peptide–water interactions, as introduced in A99-d^10^ and C36mW,^9^ and/or increased torsional barriers implemented to enforce extended peptide structures.

**Figure 1:**
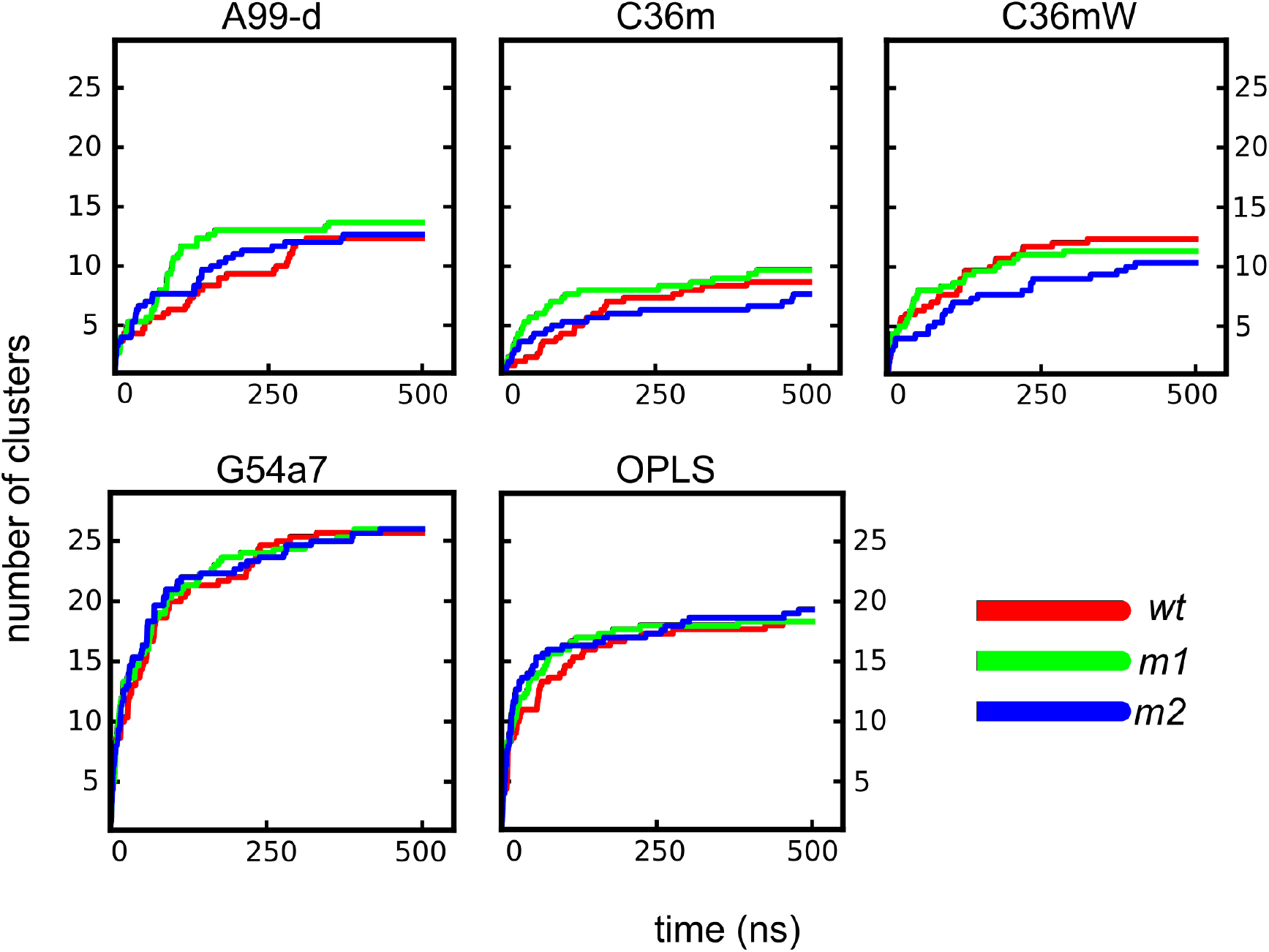
The number of structural clusters as a function of time for *wt* (red), *m*1 (green), and *m*2 (blue) from the simulations of the monomers using A99-d, C36m, C36mW, G54a7, and OPLS. Representative cluster stuctures are shown in Fig- S2.

The preference for extended structures in the simulations with the IDP-corrected FFs becomes visible by the inspection of representative structures of the most populated clusters (Fig. S2). For *wt*, A99-d, C36m, and C36mW produce an extended structure, whereas G54a7 results in a *π*-helix conformation and OPLS leads to a turn structure. A similar structural pattern is observed for *m*1 and *m*2, i.e., extended structures with A99-d and C36m(W) and structures with more intrapeptide interactions with G54a7 and OPLS. As found in previous studies we thus observe that the differences between force fields for the same peptide are larger than the differences between different peptides but using the same FF.^25^ However, it should be noted that we do not know the experimental structures of *wt, m*1 and *m*2; it is thus not clear whether they should possess different monomer structures and which of the FFs provides the better description for their conformational ensembles. If one looks into the details, some small peculiarities are found, such as a slightly reduced flexibility of *m*2 compared to *wt* and *m*1 when modeled with C36m(W), or that *m*2 is more extended than the other two peptides in the simulations with G54a7. These findings are in agreement with the higher *β*-propensity of Val compared to those of Phe and Leu. For instance, for exposed residues Fujiwara et al. assigned a *β*-propensity of 2.31, 1.50, and 1.18 for Val, Phe, and Leu, respectively.^67^

#### Intrapeptide contacts and interaction energies

To better understand the structural preferences predicted by the different FFs, we characterized the intrapeptide interactions by calculating the frequency of residue–residue contacts (Fig. 2). The resulting contacts indicate that among the FFs developed for IDPs, C36mW produces the highest preference for extended structures and A99-d the lowest. A99-d leads in particular to more contacts beyond the first- and second-neighboring residues in *m*1 and *m*2, which in the case of *m*2 is against the propensity of Val to adopt the *β*-state.^67^ C36mW is the FF that best reproduces the preference for extended structures in the order Val (*m*2) *>* Phe (*wt*) *>* Leu (*m*1). With C36m, on the other hand, this behavior cannot be reproduced, which is visible from the higher presence of intrapeptide contacts including electrostatic contact between K16 and E22 in *m*2 compared to *wt*. G54a7 and OPLS produce many intrapeptide contacts indicating collapsed monomeric structures, apart from *m*2 modeled with G54a7.

**Figure 2:**
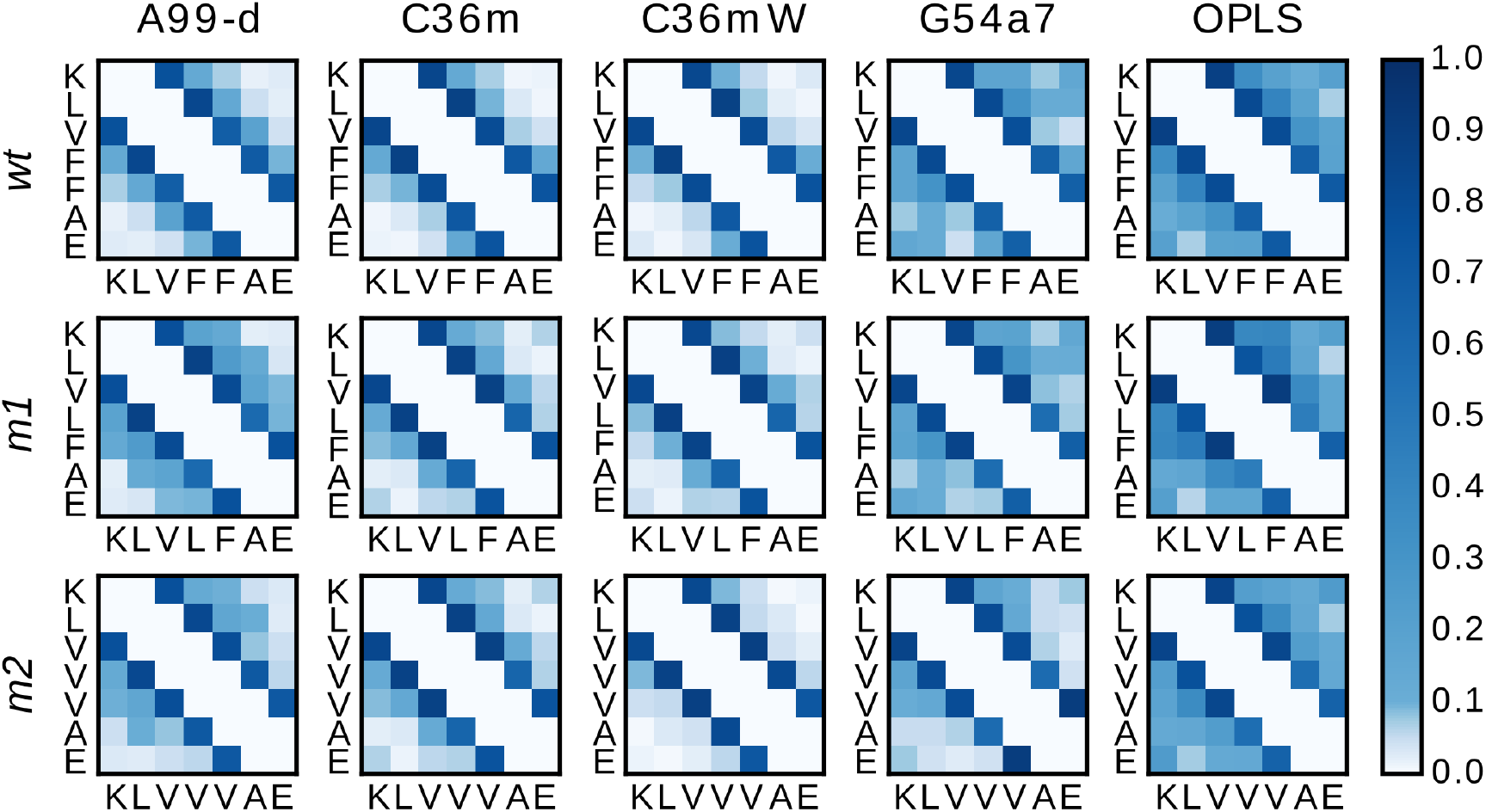
Probability of intrapeptide contacts for monomers of *wt, m*1, and *m*2 from simulations using A99-d, C36m, C36mW, G54a7, and OPLS. The color code represents the probability of a contact between residues during the MD simulations. For the sake of clarity, the diagonal and first off-diagonal elements of the contact maps corresponding to self-contacts and contacts with direct neighbors are not shown.

The intrapeptide contacts can be understood based on the inter-residue interaction energies, which can be dissected into electrostatic and vdW interactions (Fig. S3A and B, respectively). G54a7 and OPLS predict strong electrostatic interactions between K16 and E22, which cause the monomer structures to collapse, entailing several other intrapeptide contacts of vdW nature. An exception is *m*2 modeled with G54a7 where more extended structures are produced. However, here an electrostatic interaction between K16 and V18 plays a role, which is also present in *wt* and *m*1 modeled with G54a7, but to a lesser extent. This contact, which is an H-bond between the carbonyl oxygen of K16 and the amide hydrogen of V18, is only present in the simulations with G54a7 and OPLS, but none of the IDP-corrected FFs. Interestingly, with A99-d and C36m(W) the K16–E22 interaction is also of considerable intensity in *m*1, which mirrors the medium flexibility of Leu, yet does not lead to completely collapsed structures. This electrostatic attraction must thus be counteracted by solvation energies or torsional parameters of the other residues favoring overall extended structures. For *wt* and *m*1, none of A99-d, C36m, and C36mW predicts the interaction between the termini to be of (large) relevance as the corresponding interaction energies are close to zero. This suggests that these peptides are generally more extended than *m*2, which agrees with the contact maps in Fig. 2 and can be explained with the high propensity of Phe and Val to adopt a *β*-conformation. Finally, it can be concluded that for none of the FFs and peptides the vdW interactions play a large role in the structure formation of the monomer.

### 3.2 Peptide–water interactions

Peptide structure and dynamics emerge from an interplay between the peptide–peptide and peptide–solvent interactions. To understand the latter, we quantify the first hydration shell of *wt, m*1 and *m*2 and their translational diffusion.

#### Hydration shell

The first hydration shell is defined as the water layer around the peptide in which all the water molecules are directly in contact with the peptide.^68^ To fulfill this definition, one needs to i) define the peptide atoms to be used for computing the contacts, and ii) set a cutoff distance between the selected peptide atoms and the water molecules to define the contacts. There are several procedures available in the literature to obtain the cutoff distance and the peptide atoms.^69–73^ Here, we use the non-hydrogen peptide atoms to define the contacts and a single cutoff distance of *r*_cut_ = 0.45 nm between the peptide non-hydrogen atoms and the water oxygens. This particular choice emerged from the analysis of the MD trajectories using different cutoff distances and contact definitions.

Fig. S4A shows the number of water molecules in the first hydration shell on a per-residue basis. The solvation pattern is very similar across the FFs. There are only few notable differences. For *wt* we find that, except for E22, A99-d always yields slightly larger numbers of water molecules per residue than the other FFs, followed by C36mW and C36m. G54a7 and OPLS cause less solvation, especially at the C-terminal capping groups, which is accompanied by a higher K16–E22 contact probability as seen above. Thus, G54a7 and OPLS are more favorable for residue–residue interactions than the the other three FFs where the strength of the protein–water interactions was increased on purpose (A99-d and C36mW) and/or the preference for extended structures was increased. The effect of the mutations in *m*1 and *m*2 can be observed in Fig. S4B and C, respectively. For the *m*1 variant, the F19L mutation results in less hydration at the mutation site, which can be explained by the smaller volume of the side chain of Leu compared to that of Phe. For the other residues of *m*1 the differences are small. In the case of *m*2, both mutation sites F19V and F20V are less hydrated compared to *wt*, which again can be explained with the smaller side-chain size. OPLS predicts for both mutation sites of *m*2, as also for the mutation in *m*1, significantly lower differences compared to the other FFs. For the other *m*2 residues we observe that they are more hydrated compared to *wt* and also have more water molecules in the first hydration shell than the *m*1 variant. OPLS yields the largest differences for these residues. Thus, with respect to the whole peptides OPLS predicts a more similar solvation for *m*2 and also *m*1 compared to *wt*.

#### Solvent accessible surface area

One factor that determines the number of water molecules within the first hydration shell is the solvent accessible surface area (SASA) of the solute. We calculated the SASA are for *wt, m*1, and *m*2 using a standard 0.14 nm probe radius. Our results in Table S2 show that within each FF *wt* has the largest SASA and *m*2 has the lowest, which can be explained with the different sizes of Phe, Leu, and Val. The differences across the FFs for each peptide positively correlate with differences found for the number of water molecules within the first hydration shell, i.e., G54a7 and OPLS lead to smaller SASA values as a result from the more collapsed structures sampled with these two FFs. However, the interfacial water area (IWA), which is the ratio of the SASA to the number of waters within the first hydration shell, attains an almost constant value of IWA = 0.096 nm^2^ for all peptides and force fields. For globular proteins an IWA value of 0.11 nm^2^ has been previously reported^68^ suggesting that A*β*_16*−*22_ has a higher solvent density than the globular proteins.

#### Peptide–water hydrogen bonds

The analysis of the H-bonds between the peptide and water allows us to assess to what extent specific peptide–water interactions play a role, which serve as a stabilizing factor for the peptide structure and largely influence the peptide dynamics.^74,75^ To this end, we determined the number of H-bonds between peptide and water on a per-residue basis. The results of this analysis are shown for *wt* in Fig. 3. Interestingly, the propensity to form H-bonds with water is quite different from the solvation pattern found for the residues. While K16, F19, F20, and E22 are surrounded by the same amount of water molecules, E22 engages in the largest number of H-bonds, followed by K16. These two residues can use both their backbone and side chain for H-bond formation, while this is limited to the backbone for the residues L17–A21. This explains why these residues have similar numbers of H-bonds with water. We further analyzed whether the residues acted as H-bonds acceptor or donor, which is shown as dashed and non-dashed areas, respectively, in Fig. 3. The backbone can act as both a donor and an acceptor. Since the side chain of E22 can only act as an acceptor and that of K16 only as a donor, it becomes clear that most H-bonds formed by these residue take place via their side chains. While the different FFs yield the same distribution between H-bond donor and acceptor capabilities for each of the residues, G54a7 and OPLS lead to a smaller tendency for H-bond formation between peptide and water. This difference is most pronounced for E22, even though OPLS led to the same level of hydration for that residue.

**Figure 3:**
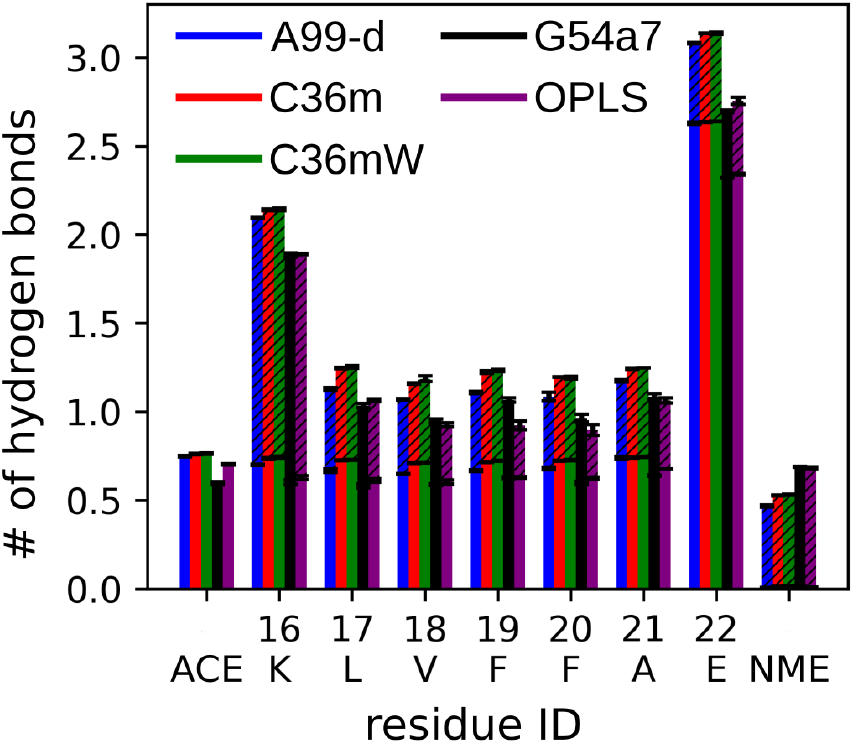
The average number of H-bonds formed between *wt* residues and water. The dashed and non-dashed areas indicate the number of instances where the amino acids act as H-bond donor and acceptor, respectively. The color key for the FFs is given as part of the plot.

#### Translational diffusion constants

The three-dimensional translational diffusion constants for *wt, m*1, and *m*2 are shown in Table 1, which were calculated from the MSDs (Fig. S5). The experimental value for the wild-type A*β*_16*−*22_ has previously been measured by NMR spectroscopy and has a value of 0.353*×*10^*−*5^ cm^2^/s.^76^ Our results show that none of the FFs can reproduce the experimental three-dimensional translational diffusion constant. A99-d yields considerably smaller diffusion constants than observed in experiment, whereas the other FFs overestimate the diffusion, with C36m leading to the fastest diffusion. However, after correcting the diffusion constants by the viscosity of the solvent models,^77^ all FFs underestimate the diffusion similarly to A99-d. While there are small differences between the computed diffusion constants for *wt, m*1, and *m*2, most of them obtained per FF can be considered identical within the standard error. Exceptions are the diffusion constants resulting from the C36m(W) simulations. However, here no clear trend is observed: with C36m the mutant *m*2 has the smallest diffusion constant, whereas with C36mW the wild-type peptide has a smaller diffusion constant than its mutants. Since we have no experimental results for the translational diffusion constants for *m*1 and *m*2, we are unable to judge the latter findings.

**Table 1:** Three-dimensional translational diffusion constant *D*_*t*_. The values in parentheses are *D*_*t*_ scaled by the correction factor of the solvent model viscosity (where available). The scaling factors are 2.8 for TIP3P (C36m), 3.27 for SPC (G54a7), and 2.13 for TIP4P (OPLS).^77^

|  | A99-d | C36m | C36mW | G54a7 | OPLS |
| --- | --- | --- | --- | --- | --- |
| *wt* | $0.21 \pm 0.01$ | $0.69 \pm 0.04$<br>(0.25±0.02) | $0.56 \pm 0.01$ | $0.52 \pm 0.05$<br>(0.16±0.02) | $0.47 \pm 0.03$<br>(0.22±0.01) |
| *m1* | $0.21 \pm 0.01$ | $0.66 \pm 0.02$<br>(0.24±0.01) | $0.61 \pm 0.06$ | $0.54 \pm 0.05$<br>(0.17±0.02) | $0.46 \pm 0.04$<br>(0.22±0.02) |
| *m2* | $0.21 \pm 0.01$ | $0.59 \pm 0.05$<br>(0.21±0.02) | $0.62 \pm 0.02$ | $0.53 \pm 0.04$<br>(0.16±0.02) | $0.43 \pm 0.04$<br>(0.20 ±0.02) |

### 3.3 Characterization of the aggregation process

#### Oligomer size

After studying the conformations of single peptides in solution, we turn our attention to the aggregation properties of A*β*_16*−*22_ and its mutants represented by the different FFs. We choose to study the aggregation properties by simulating six peptides in a water box. Fig. 4 shows the size of the oligomers as a function of time. We observe that the peptides aggregated more quickly attaining the highest oligomer size within 300 ns in the case of G54a7 and OPLS as compared to A99-d, C36m, and C36mW. Despite the intrapeptide interactions observed in the monomer state being of similar strength for A99-d and C36m(W), the highest average oligomer size achieved with A99-d was a trimer for *wt* and a dimer for the variants *m*1 and *m*2. With C36m(W), on the other hand, the highest oligomer size is reached for all peptide systems, albeit for *m*2 the oligomers are less stable. Both *wt* and *m*1 aggregate more quickly than *m*2, while the former two peptides aggregate with a similar speed. C36mW predicts the smallest aggregation tendency for *m*2. Thus, from the five FFs under study, C36m and C36mW are the only ones which are able to model *m*2 as less aggregation-prone than *wt* and *m*1 as found in experiments.^27^ However, the correct behavior according to experiment would be that *m*2 does not aggregate at all, while *m*1 aggregates faster than *wt*. This aggregation ranking cannot be modeled by any of the five FFs and also none of the FFs that were included in our previous benchmark. ^12^ While G54a7 and OPLS overestimate the aggregation tendency of all three peptides,^12^ A99-d underestimates it.

**Figure 4:**
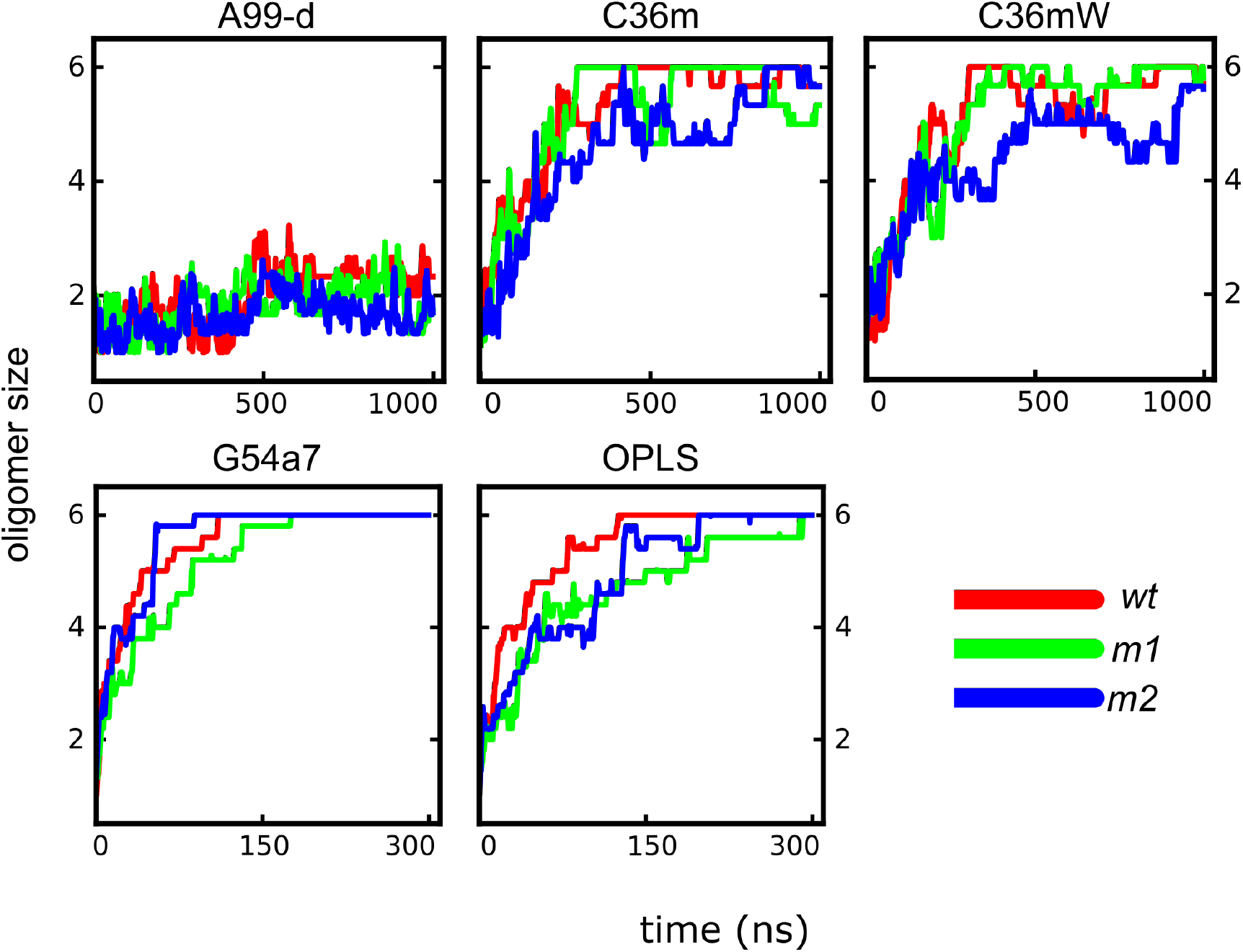
Average oligomer size as a function of time for *wt* (red), *m*1 (green), and *m*2 (blue) using A99-d, C36m, C36mW, G54a7, and OPLS. The graphs are averages over three trajectories per system in the case of A99-d and C36m(W) and five trajectories for G54a7 and OPLS. The different *x*-axes scales for the top and bottom row are noted.

#### Transition networks and oligomer structures

For the characterization of the intermediate oligomeric states and the transitions between them, we calculated transition networks (TNs).^34,58^ As in our preceding study,^12^ we chose to build the TNs in a two-dimensional space defined by *β*-strand content (*x*-axis) and oligomer size (*y*-axis) (Fig. 5). The TNs for A99-d confirm that this FF does not support stable oligomer formation; for all three peptides the most populated state is the extended monomer structure. All oligomer states are only weakly populated with the maximum oligomer size being a pentamer in the case of *wt* and tetramers for *m*1 and *m*2. Typical snapshots for oligomers formed with A99-d (Fig. 6) show that the peptides in these oligomers are only loosely attached and no *β*-sheets are formed. As we used the backbone dihedral angles for the definition of the *β*-state and since with A99-d the peptides have a high preference for extended structures, these oligomers nonetheless populate TN states with high a *β*-content. In fact, the three peptides modeled with A99-d only aggregate when being extended, which is understandable as in the collapsed peptide state, K16 and E22 interact with each other within the peptide as seen from the monomer simulations.

**Figure 5:**
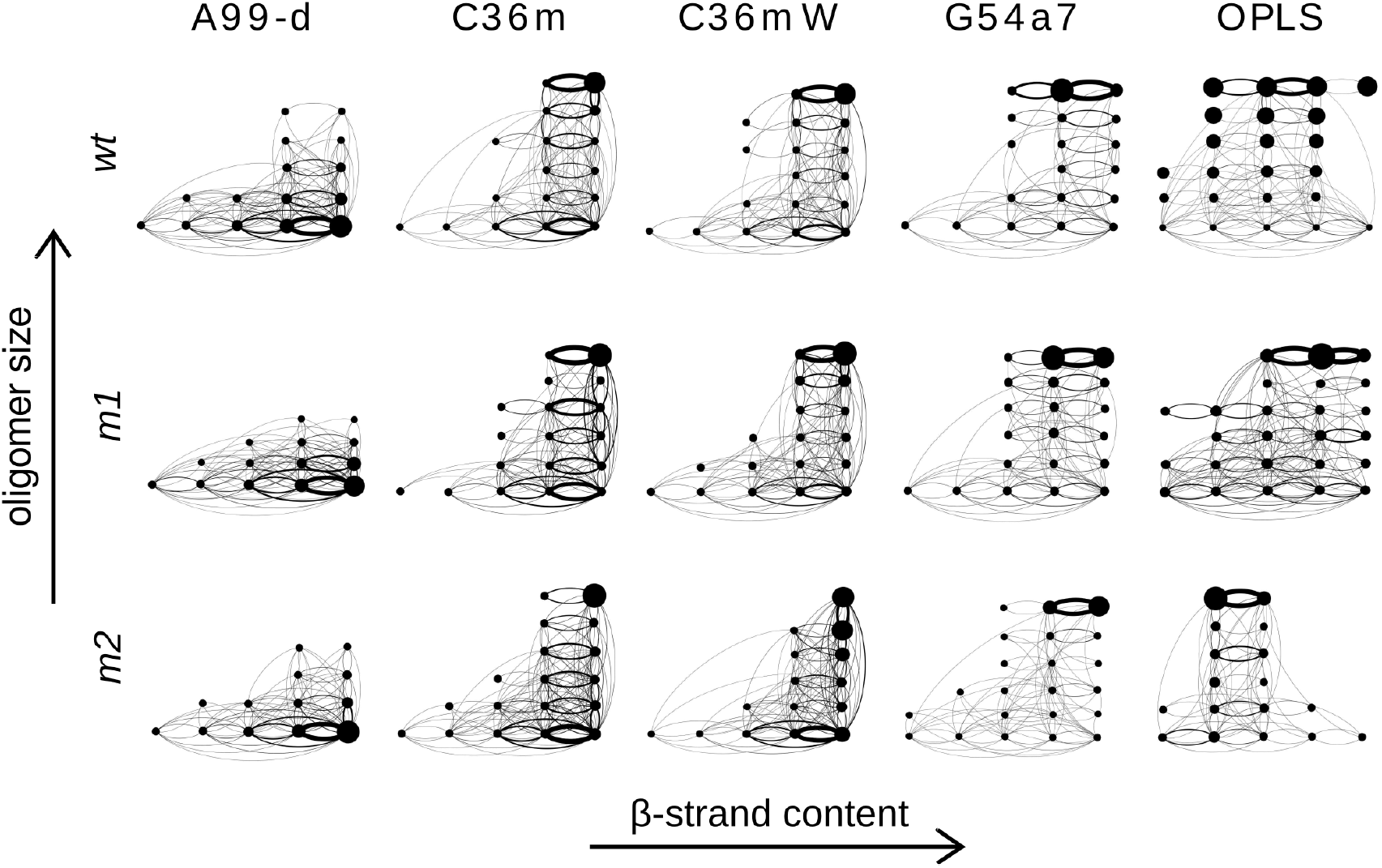
Transition networks for the aggregation of *wt* (top), *m*1 (middle), and *m*2 (bottom) using A99-d, C36m, C36mW, G54a7, and OPLS. The oligomer size (from monomer to hexamer) is given along the vertical axis and the horizontal axis represents the *β*-strand content (divided into 5 ranges: 0–20%, 20–40% etc.). The size of the nodes is proportional to the population of the state, and the width of the edges is proportional to the mass flux between the states.

**Figure 6:**
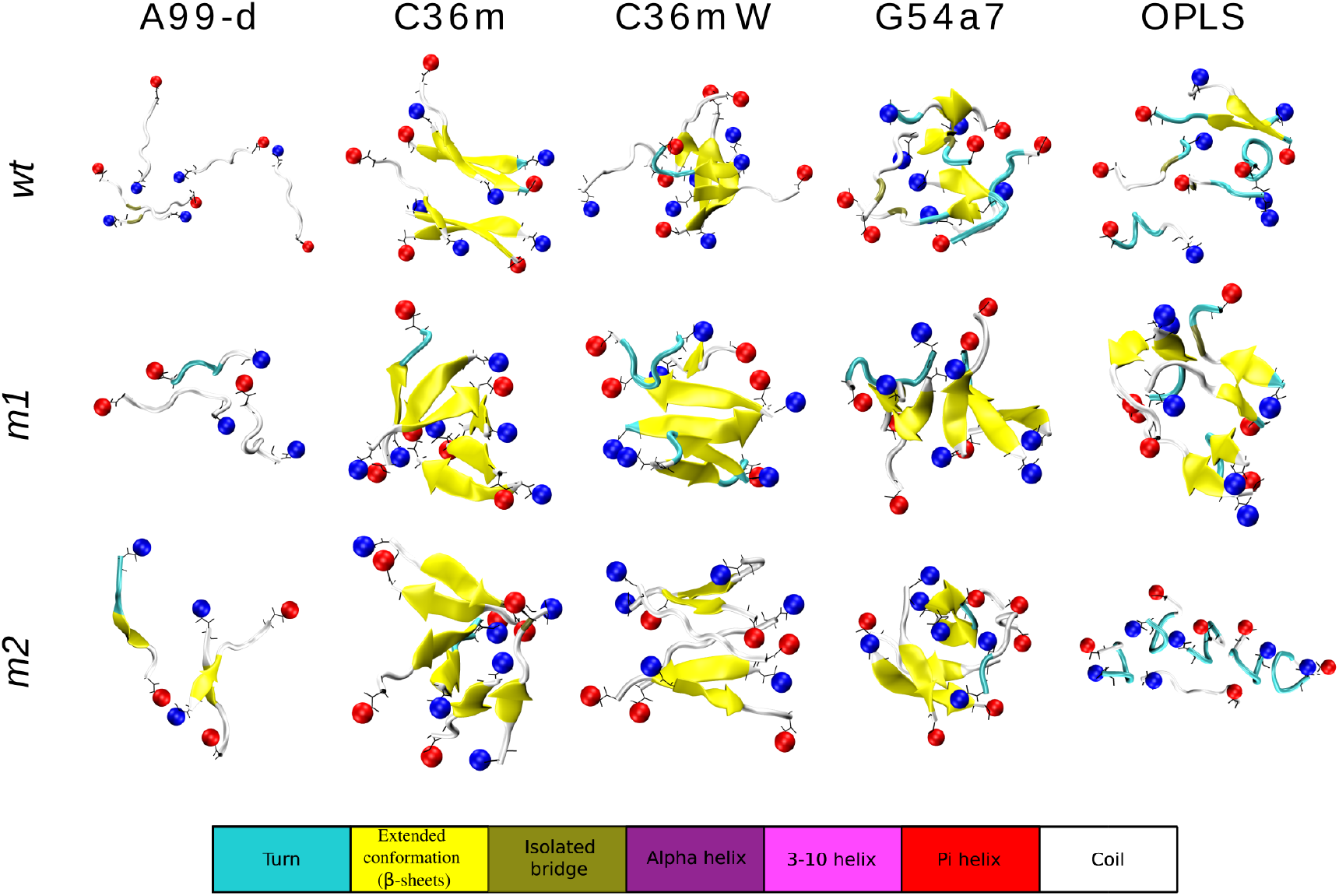
Representative structures of the largest oligomers formed during the MD simulations of oligomer formation of *wt* (top), *m*1 (middle), and *m*2 (bottom) modeled with A99-d, C36m, C36mW, G54a7 and OPLS. The spheres colored in red and blue represent the N- and C-terminus, respectively. The assignment of the secondary structure is according to the color key at the bottom.

The most populated state in the TNs obtained with C36m and C36mW is in each case a hexamer with a high amount of *β*-sheets. For *m*2 modeled with C36mW the pentameric state is of almost equal importance, corroborating the finding that with this FF the aggregation is slowed down for this peptide. Typical structures indicate that with C36m(W) well-developed *β*-sheets are adopted. They are characterized by a high amount of antiparallel *β*-sheets resulting from the electrostatic attraction between neighbored K16 and E22 within a sheet. In some cases two sheets with three peptides each form a steric zipper as best seen for *wt* modeled with C36m in Fig. 6. Such a double layer can serve as a nucleus for amyloid fibril formation. In other cases, such as *m*2 modeled with C36m barrel formation is observed. With both C36m and C36mW the aggregation pathways involve *β*-sheet formation from the very beginning, i.e., no disordered aggregates with a low amounts of *β*-content are formed. This behavior can be understood based on the high tendency of the peptide monomers to adopt extended structures. While the latter also holds true for A99-d, C36m and C36mW provide a better balance between peptide–peptide and peptide–water interactions.

As already seen in our previous study,^12^ G54a7 and OPLS result in the formation of disordered oligomers with lower amounts of *β*-sheets, which mirrors the behavior of the monomers when modeled with these two FFs. The most extreme case is given by *m*2 modeled with OPLS. As the representative snapshot of the hexameric state of this system shows, here the peptides mainly adopt a turn conformation as a result of strong intrapeptide K16–E22 attraction (see Fig. S2A), inhibiting *β*-sheet formation in this system. Compared to OPLS, G54a7 provides a better description of the amyloid oligomers formed by A*β*_16*−*22_ and its mutants, especially for the largest oligomer considered here, which, after it formed, relaxed into *β*-sheet conformations. However, compared to the structures found for C36m(W) these *β*-sheets are less ordered.

#### Interpeptide contacts and interaction energies

To better understand the driving forces behind the different aggregation patterns observed for the different FFs, we first analyzed the interpeptide contacts present in the oligomers that formed (Fig. S6). However, we find that these contacts are too unspecific to allow for an in-depth understanding. Instead, we turned our attention to the nonbonded interaction energies between the peptides on a per residue–residue basis. The resulting energy decomposition is shown for the vdW and electrostatic energy contributions in Fig. 7 and Fig. 8, respectively, while the sum over all residue–residue interaction energies per system are provided in Table S3.

**Figure 7:**
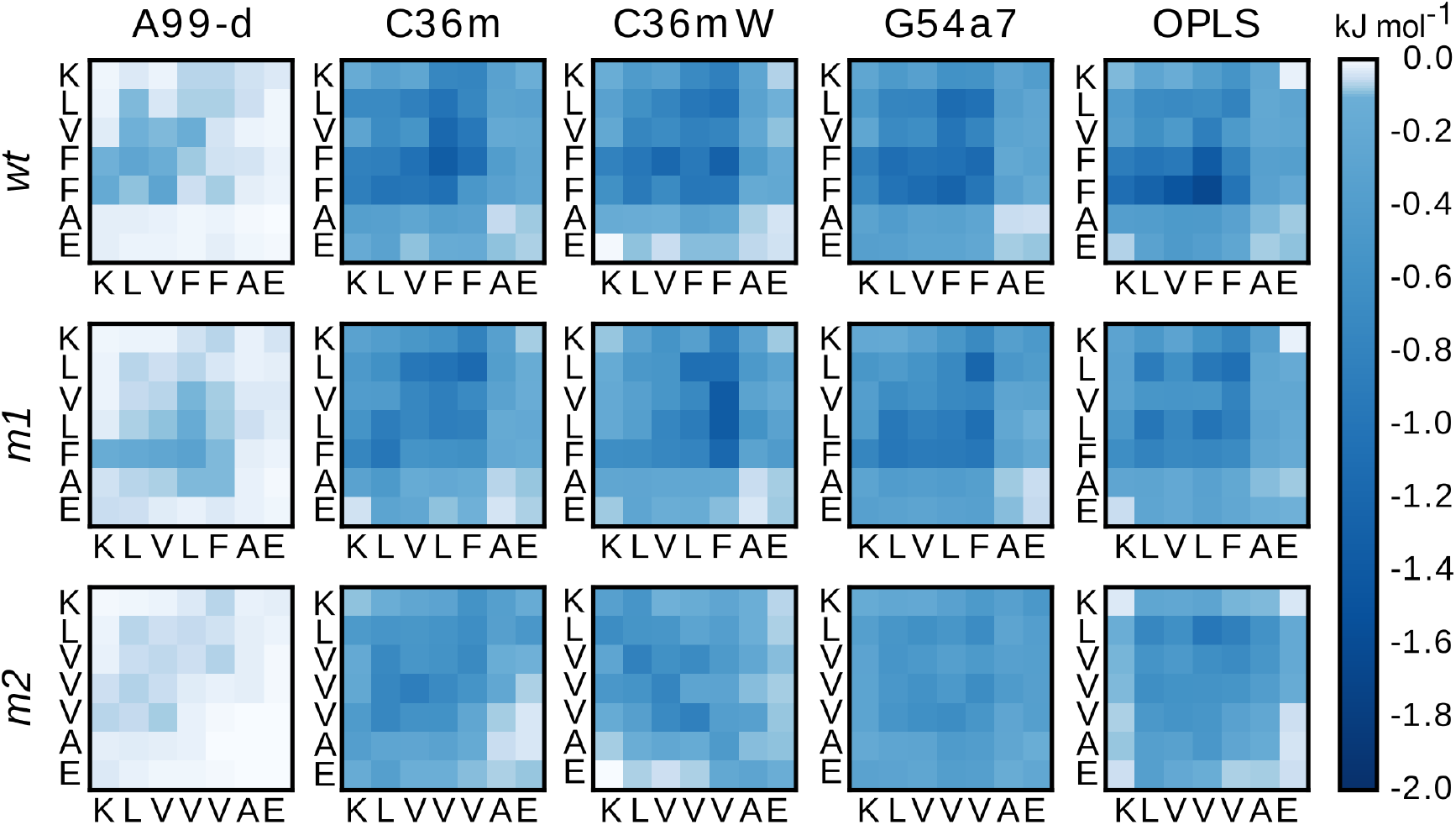
Residue–residue vdW interaction energies between peptides forming oligomers of *wt* (top), *m*1 (middle), and *m*2 (bottom) obtained from simulations using A99-d, C36m, C36mW, G54a7, and OPLS. The interaction energies (in kJ mol^*−*1^) are according to the color key on the right.

**Figure 8:**
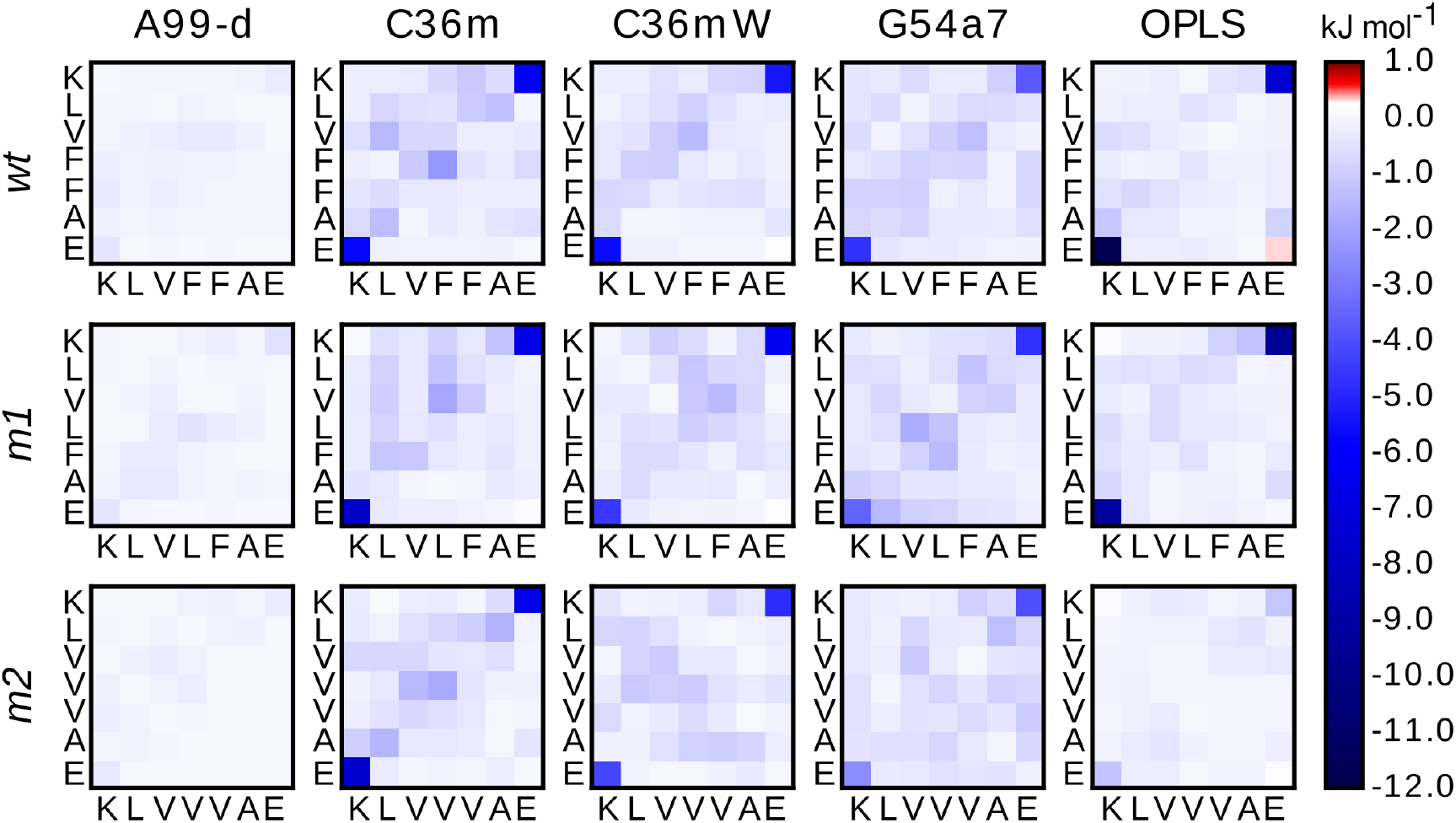
Residue–residue electrostatic interaction energies between peptides forming oligomers of *wt* (top), *m*1 (middle), and *m*2 (bottom) obtained from simulations using A99-d, C36m, C36mW, G54a7, and OPLS. The interaction energies (in kJ mol^*−*1^) are according to the color key on the right.

For A99-d only weak interaction energies are seen in the interaction maps, and the values in Table S3 show that the level of interaction is an order of magnitude weaker than with the other FFs. This explains why with A99-d the peptides did not aggregate: the peptide–water interactions are stronger than the interactions between the peptides. With the other FFs strong vdW interactions are observed among residues from the hydrophobic core regions of *wt* and *m*1, while for *m*2 they are generally weaker, which is confirmed by the accumulated values in Table S3. For this peptide also the electrostatic interactions are weaker with each of these FFs. In principle, this could translate into a reduced aggregation propensity as seen experimentally,^27^ which however is not the case, especially not with G54a7 and OPLS. In particular with OPLS the interpeptide interactions for *m*2 are considerably weaker than those for *wt* and *m*1, yet the aggregation speed is largely the same for all three peptides. Thus, for OPLS the peptide–water interactions must be too weak so that the peptide–peptide interactions, even when not as pronounced, dominate the aggregation behavior. The analysis of the hydration shell around monomers indeed revealed that solvation is of less importance in G54a7 and OPLS. With C36mW, on the other hand, the increased protein–water interactions result in somewhat weaker interactions between the peptides, yet opposite to A99-d they are still strong enough to enable peptide aggregation.

In general, all FFs predict very similar total interpeptide interaction energies for *wt* and *m*1, which explains why both peptides aggregated at similar speeds in the MD simulations. The contribution of the electrostatic interactions to the total interaction potentials is in most cases slightly larger than the vdW contribution. Exceptions are *wt* and *m*1 modeled with A99-d and *m*2 simulated with OPLS. The dominant electrostatic interaction is the K16–E22 contact, causing the peptides to arrange themselves in an anti-parallel orientation to each other, which was best seen in the simulations with C36m(W). As mentioned above, *m*2 simulated with OPLS prefers to form a turn structure with intrapeptide K16–E22 salt bridges, which prevents these two residues from interacting with each other across peptides.

### 3.4 Stability of preformed fibrillar aggregates

In the previous section, we characterized the self-assembly of six randomly positioned monomers of *wt, m*1, and *m*2. Here we test whether the various FFs are able to maintain the structure of the aggregation end product, which is a fibril. If this is not possible it follows that one can also not simulate the aggregation into that state. To this end, we set up a minifibril involving two *β*-sheet layers with six peptides forming an in-register, antiparallel *β*-sheet in each layer (Fig. S1). The stability of this minifibril, also called steric zipper,^54^ was tested with 3 *×* 300-ns MD simulations per system using the recently developed force fields A99-d, C36m, and C36mW. We performed various analyses to test the stability of the simulated minifibril. One of them is the RMSD (Fig. S7), which shows that the systems are less stable when simulated with A99-d. The largest RMSD is observed for *wt* modeled with A99-d, which results from one of the twelve peptides leaving the minifibril at *t ≈* 200 ns (Fig. S8). However, this occurred during only one of the three simulations for this system. From the RMSD plots for C36m and C36mW one can see that with these two FFs the *wt* minifibril is most stable, while it is least stable for *m*2.

In order to distinguish whether the RMSD changes arise from a slow disassembly of the fibril structure or from local fluctuations, we calculated the nematic order parameter *S*_2_ defined in eq (1) to describe the orientational order of the peptides with respect to each other (Fig. 9). This analysis clearly shows that with A99-d the characteristic arrangement of peptides within a fibril is not supported with A99-d. For all three peptide sequences *S*_2_ drops below 0.5, for *m*1 and *m*2 it even reaches values as as low as *≈*0.3. The loss of the steric zipper geometry can also be seen in the representative structure shown in Fig. S8. With C36m and C36mW the overall fibril structure is generally better retained, especially for *wt* modeled with either FF where *S*_2_ *≈* 0.8 at the and of the 300-ns simulations. This agrees to the findings of the hexamer simulations, which already revealed steric zipper formation with three peptides per sheet for *wt* (Fig. 6). The double mutant *m*2, on the other hand, appears not to be stable in the fibril state as *S*_2_;S 0.6 at the end of the simulations. The hexamer simulations for *m*2 revealed a slowed-down aggregation, yet *β*-sheets were nonetheless formed. The combined picture from both sets of simulations thus suggests that C36m and C36mW model a reticent aggregation of *m*2 into *β*-sheets, which however would not develop into a fibril. This only partly agrees with the experimental findings,^27^ based on which no aggregation at all should be happening. The fibril structure of *m*1 is found to be stable when modeled with C36mW, yet starts to disintegrate when simulated with C36m. Thus, C36mW is better suited to reproduce the intricacies of the aggregation of A*β*_16*−*22_ and its mutants.

**Figure 9:**
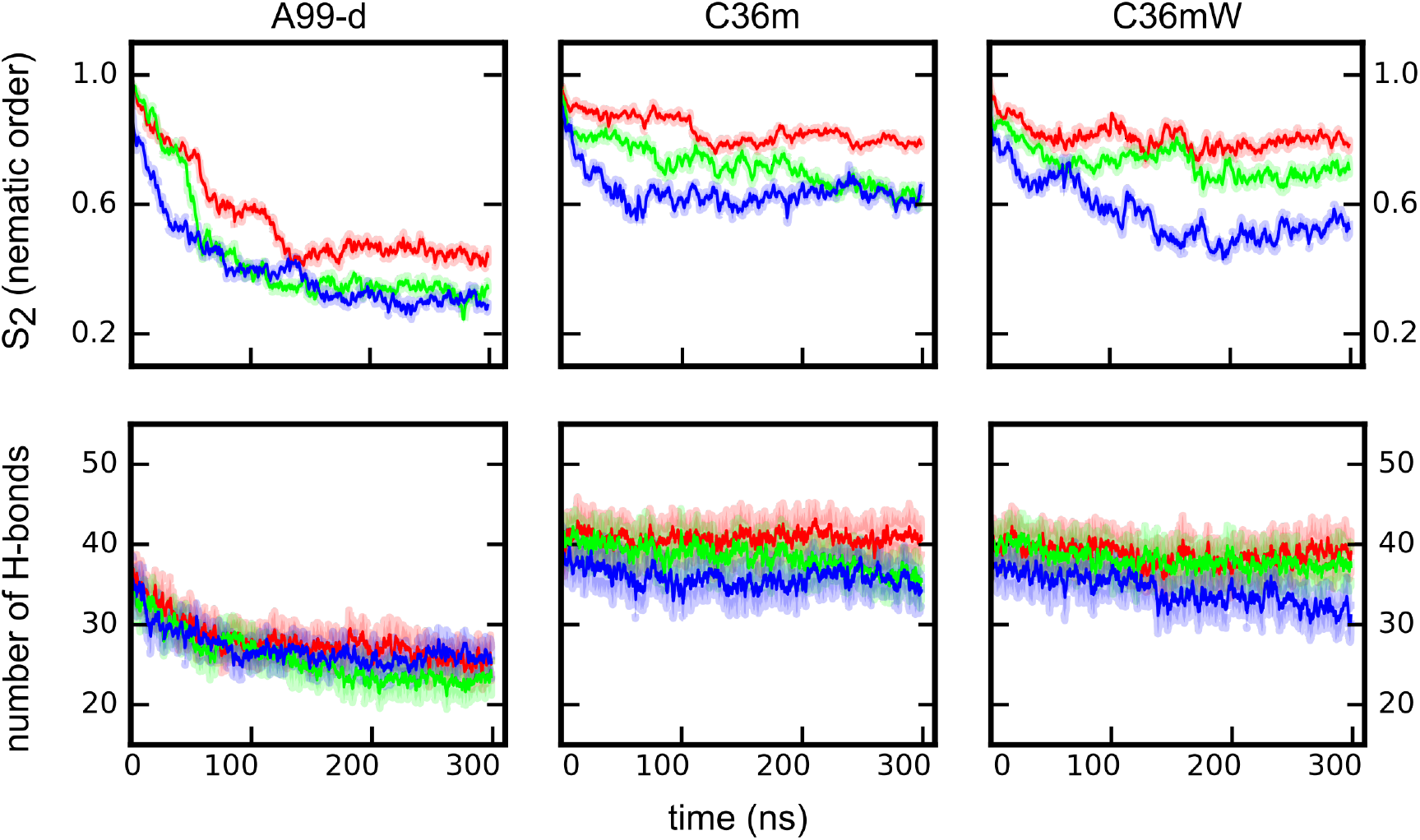
The change in the nematic order parameter (top) and the number of backbone H-bonds (bottom) for the minifibril of *wt* (red), *m*1 (green), and *m*2 (blue) simulated with A99-d, C36m, and C36mW. The averages over three independent simulations per system are shown. The shaded areas indicate the standard error.

To gain further insight into the origin of the fibril instabilities seen for some of the systems, we determined the *β*-sheet content as well as the number of H-bonds formed between peptides within the two *β*-sheets. In Fig. 9 the average number of H-bonds per sheet is shown. The maximum number is *≈*40, which includes both backbone and side-chain H-bonds. In the simulations with A99-d the number of H-bonds consistently decreased, reaching values between 20 and 30 for all three peptides. The breaking of H-bonds is accompanied by a dissolution of *β*-sheets, as Fig. S7 shows, yet the decline is not as pronounced as for the H-bonds. Taken together, the fibril simulations confirm that with A99-d peptide aggregation cannot be simulated as the peptide–water interactions are considerably more favorable than interpeptide H-bonds. With C36m(W), on the other hand, the H-bonds and *β*-sheets are well conserved for *wt* and with C36mW also for *m*1, while for *m*2 the number of H-bonds has dropped to 30–35 at the end of the 300-ns MD simulations. The count of H-bonds for *m*1 modeled with C36m remained constant until 200 ns after which it started to decline somewhat. Thus, this analysis confirms that from the FFs considered C36mW is best suited for modeling *wt, m*1, and *m*2, followed by C36m.

## 4 Discussion and conclusions

In this work, we examined the applicability of current force fields (FFs) developed for IDPs, namely AMBER99SB-*disp*^10^ *(A99-d), CHARMM36*m^9^ (C36m), and C36m with increased protein–water interactions^9^ (C36mW), for studying amyloid aggregation with MD simulations using the A*β*_16*−*22_ peptide and its two mutants F19L and F19V/F20V (denoted *wt, m*1, and *m*2 here) as test case. Based on experimental results the order of the aggregation speed should be *m*1 *> wt » m*2 *≈* 0.^27^ As amyloid aggregation results from an interplay of monomeric peptide properties as well as the stability of the intermediate oligomers and the final aggregation product, which are amyloid fibrils, we employed a step-wise approach and investigated each of these aspects separately. In order to gain an in-depth understanding of the performance of the FFs we dissected the intra- and interpeptide interaction energies of the monomers and oligomers, and included in our analysis the results of the simulations obtained with GROMOS54a7 (G54a7) and OPLS-AA (OPLS). For these two FFs we had already demonstrated that they are not suitable for the problem under study, because they overestimate the aggregation process and cannot discriminate between the different aggregation propensities of *wt, m*1, and *m*2.^12^ By including them in the analysis we wished to learn the origin of their failure, knowledge that in future can be used for FF reparameterization. For the IDP-corrected FFs we can conclude that A99-d is not applicable to the process of amyloid aggregation as this FF does not lead to stable aggregates being formed, whereas C36m(W) give rise to proper *β*-sheet formation with the aggregation speed order *wt* ≳ *m*1 *> m*2. In the following we discuss how these results are affected by the properties at the monomeric and oligomeric level as well as how the components of the FFs give rise to their different behavior.

### Different peptide structures with different force fields

Given the quite small size of the peptides under study, one would expect that the different FFs should not lead to large differences in the peptide structures and dynamics sampled in the MD simulations. However, Figs. 1 and 2 show that this is not the case. The older FFs, G54a7 and OPLS, lead to a considerably larger structural diversity than the IPD-corrected FFs. Nonetheless, this high flexibility at the monomeric level does not hinder the peptides from rapid self-association as can be seen in Fig. 4. The larger structural diversity with G54a7 and OPLS is also visible at the oligomeric level, where especially for the smaller oligomers structures other than *β*-sheets are also sampled. This can be deduced from the transition networks in Fig. 5. In the simulations with A99-d and C36m(W), on the other hand, extended structures prevail both at the monomeric and oligomeric level. This leads to ordered aggregation into *β*-sheets from the very beginning in the case of C36m(W), whereas with A99-d only encounter complexes that promptly dissociate again are formed.

### Peptide–water interactions as key determinant for peptide structure and aggregation

Our analysis of the intra- and interpeptide interaction energies as well as the interactions between peptide and water revealed that the preference of A*β*_16*−*22_ and its mutants to adopt collapsed structures at the monomeric level and disordered oligomer structures when modeled with G54a7 and OPLS, but extended structures when modeled with A99-d and C36m(W) is mostly due to the different amounts of peptide–water interactions. Fewer H-bonds between the peptides and water are formed in the simulations with G54a7 and OPLS, which can be seen in Fig. 3. This allows for more intra- and interpeptide interactions being formed, which explains both the larger structural diversity as well as the faster aggregation. The IDP-corrected force fields, on the other hand, lead to extended structures as here especially the terminal residues K16 and E22 have a high preference to interact with water via H-bond formation, inhibiting their interaction with each other which would lead to a collapsed peptide structure. However, in the case of A99-d the peptide–water interactions are too much increased, leading to reduced interpeptide interactions and thus inhibiting aggregation. A better balance between peptide–water and peptide–peptide interactions is reached with the other FF with increased vdW interactions between protein and water, i.e., C36mW. While for the remaining FFs the overall strength of the residue–residue interactions is similar, specific interactions nonetheless play a role in the structural preferences at the monomer and oligomer level. For instance with OPLS the electrostatic attraction between K16 and E22 is very strong, causing turn structures in the monomers and fast, electrostatic-driven aggregation.

### Too strong peptide–water interactions with A99-d

In the case of A99-d not only the number, but also the strength of the H-bonds between peptide and water is increased. This can be deduced from the much smaller translational diffusion constant obtained with this FF compared to the four other ones (Table 1). We conclude that this is the main source of the decreased aggregation propensity observed with A99-d. A similar behavior was seen in the simulations with AMBER03WS in our previous study,^12^ a FF where the short-range protein–water pair interactions were increased by a factor of 1.1, while all the water–water and protein–protein parameters were left unchanged.^7^ However, it should be mentioned that also the peptide conformation may play a role in the simulations with A99-d. While the peptides prefer extended structures, these are in fact to a large extent poly-proline II (PPII) conformations. This shortcoming of A99-d was already revealed in our previous work^78^ and confirmed for A*β*_16*−*22_ here (data not shown). It has been proposed that PPII might prevent the formation of *β*-sheet^79^ due to its particular orientation of the amide bonds. ^80^ Thus, the problems of A99-d are twofold and involve both backbone and protein–water interaction parameters.

### Best performance for C36mW

A generally better performance was obtained with C36m and C36mW. The differences between their results are subtle. While the numbers of H-bonds between peptides and water are identical, with C36mW they are somewhat stronger – judged by the translational diffusion – as a result of increasing the protein–water interactions. ^9^ This parameterization change slightly increased the solvation of *m*2 modeled with C36mW compared to C36m (Fig. S4), which leads to a decreased aggregation propensity as both the hexamer and the minifibril simulations showed. Thus, only small adaptations of of the protein–water interactions as done in C36mW^9^ can make a positive difference. Nonetheless, the results obtained with C36 and C36mW are rather similar, allowing to conclude that C36m is in general suitable for modeling peptide aggregation. This is partly due to the refined backbone torsion parameters leading to more extended peptide structures,^9^ but also seems to be a general feature of CHARMM force fields since in our previous study CHARMM22* was already identified as the best FF for modeling the aggregation of *wt, m*1, and *m*2. Also in our benchmark of A*β*_1*−*40_ the performances of C36m and CHARMM22* were quite similar.^78^

### Future directions

The overall conclusion is that the IDP-corrected FFs A99-d and C36m(W) provide no or only minor advancements to previous FFs when it comes to modeling peptide aggregation. Neither the older^12^ nor the newer FFs can reproduce the different aggregation propensities of A*β*_16*−*22_ and its two mutants, F19L and F19V/F20V. Moreover, also all FFs considered here failed to predict the experimental diffusion constant of A*β*_16*−*22_. Thus, further FF parameterization is required. This conclusion considers the results of applying AMBER99SB-ILDN/TIP4P-D to the aggregation of *wt, m*1, and *m*2. Also with this water model, which implies increased water dispersion interactions compared to TIP4P^8^ and which were further increased in A99-d,^10^ no stable peptide oligomers formed. This result reinforces our conclusion that a general and quite drastic increase of the peptide–water interactions is not the right way to go. Instead, we propose to carefully reparameterize the Lennard-Jones interactions between protein and water taking the characteristics of each amino acid into account, with the aim to reproduce the free energy of solvation for individual amino acids as well small peptides. In addition, any future (re)parameterization attempt should aim at reproducing further experimental observables, such as different aggregation propensities as available for A*β*_16*−*22_ and it mutants,^27^ the translational diffusion of different peptides, as well as conformational characteristics of the monomeric state, given that the current study showed that the different behaviors of the FFs already emerge for the peptide monomers. Only when a diverse set of target properties is included during FF development, one will have a chance to optimize the large set of FF parameters without introducing dominant biases, as seen here for A99-d favoring the solvated protein state or with G54a7 and OPLS targeting the folded state. For the time being, the recommendation is to use C36mW; while it is not a perfect FF either, overall it produced the best results for the current aggregation benchmark and is one of the better FFs when it comes to modeling full-length A*β* ^78^ and other IDPs.^9,10^

## Acknowledgements

The authors thank Dr. Martín Carballo-Pacheco for fruitful discussions. Simulations were performed with computing resources granted from RWTH Aachen University on CLAIX-18 cluster under project rwth0400.

## Supporting information available

Figures showing the initial structure of the minifibril, representative structures for the monomers of *wt, m*1, and *m*2, intrapeptide interaction energies for the monomers, the number of water molecules around the residues of monomeric *wt, m*1, and *m*2, the mean squared displacement of the monomeric peptides, interpeptide contacts in oligomers, the RMSD and *β*-sheet content during the simulations of the minifibril, a representative structure showing peptide detachment from the fibril when simulated with A99-d. Tables listing the populations of the first two structural clusters obtained for the monomers, their SASA and hydration levels, and cumulative interpeptide interaction energies in oligomers.

## Graphical TOC Entry

**Figure 10:**
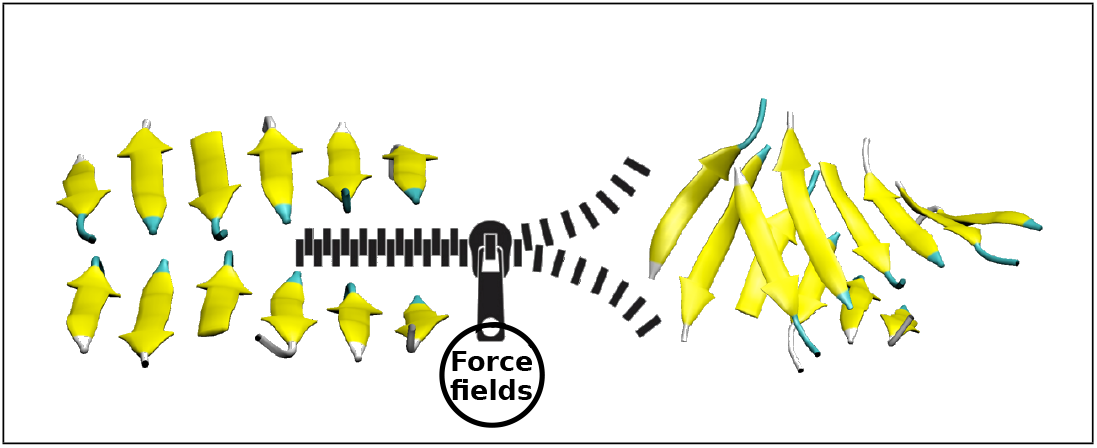
TOC graphic

## Notes

### Competing Interest Statement

The authors have declared no competing interest.

