## Supporting Information for "Different force fields give rise to different amyloid aggregation pathways in molecular dynamics simulations"

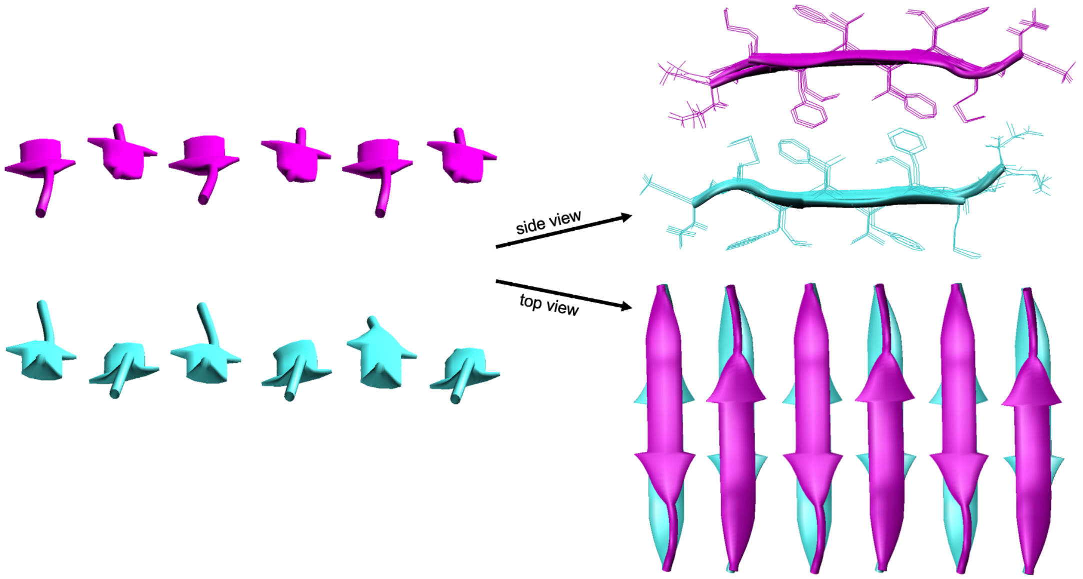


Figure S1: The initial structure of the minifibril.

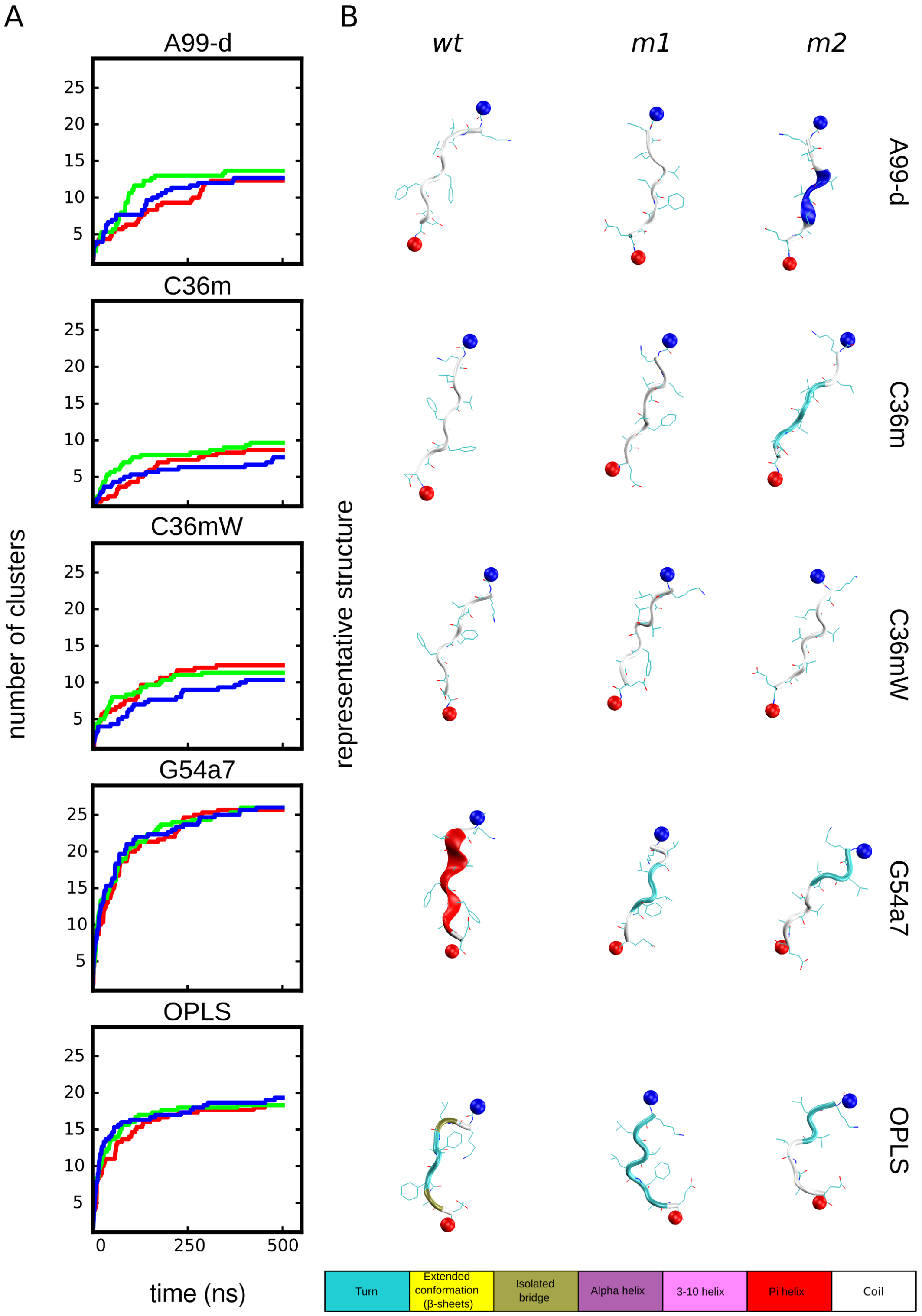


Figure S2: (A) The number of clusters as a function of time for *wt* (red), *m*1 (green), and *m*2 (blue) from the simulations of the monomers using A99-d, C36m, C36mW, G54a7, and OPLS. (B) Representative peptide structures from the corresponding most populated clusters. The spheres colored in red and blue represent the N- and C-terminus, respectively. The assignment of the secondary structure is according to the color key at the bottom.

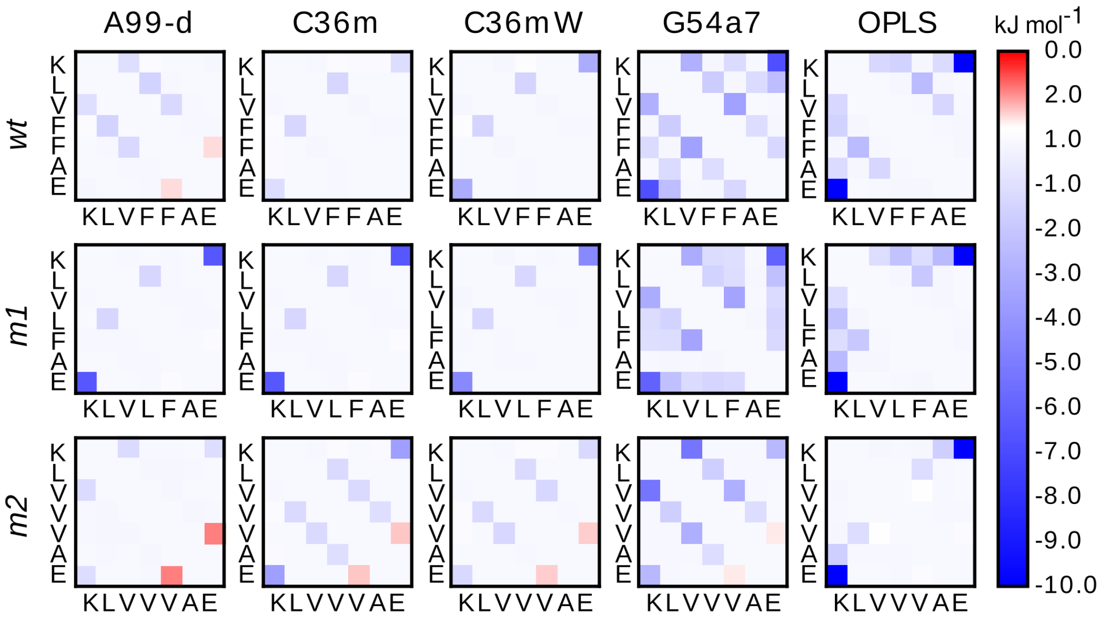


A

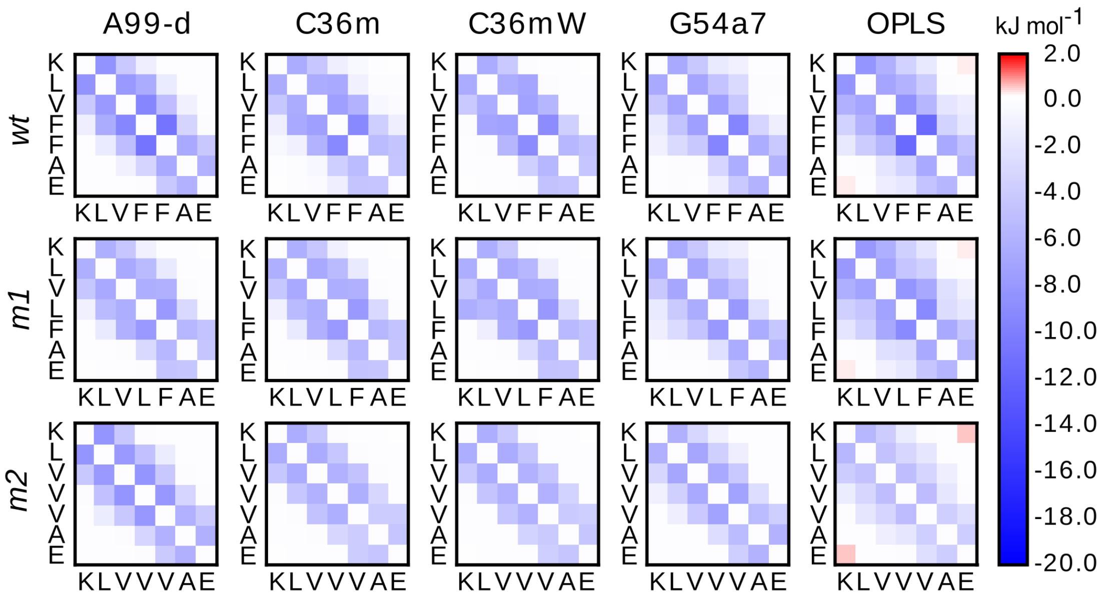


B

Figure S3: Intrapeptide electrostatic (A) and vdW (B) interaction energies for monomers of *wt*, *m*1, and *m*2 from simulations using A99-d, C36m, C36mW, G54a7, and OPLS. The colors represent the energies according to the scale on the right. The red and blue colors denote repulsive and attractive interactions, respectively. For the sake of clarity, the diagonal and first off-diagonal elements of the interaction energies corresponding to interactions with first and second neighbors in the sequences are not shown.

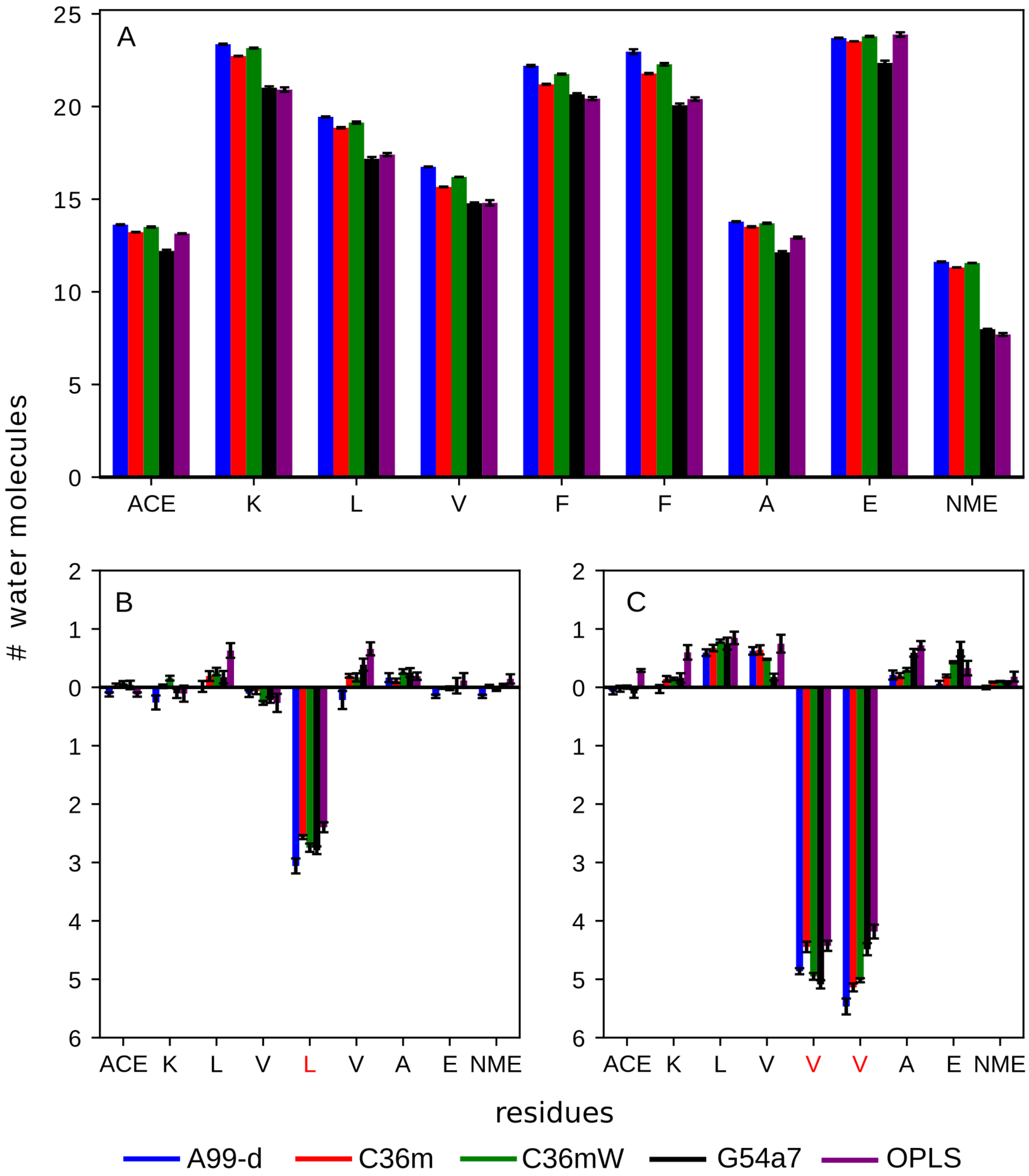


Figure S4: (A) The number of water molecules at a cutoff distance $r_{\text{cut}} = 0.45$ nm around each residue for the *wt* peptide. (B) and (C) The difference between the number of water molecules at a cutoff distance $r_{\text{cut}} = 0.45$ nm per residue for $m1-wt$ and $m2-wt$, respectively. The amino acid sequence for each peptide is given at the $x$-axis. The red-labeled residue names correspond to the mutations in (C) and (D). Values averaged over three trajectories are shown, along with the standard error of the mean. The color key for the FFs is given at the bottom.

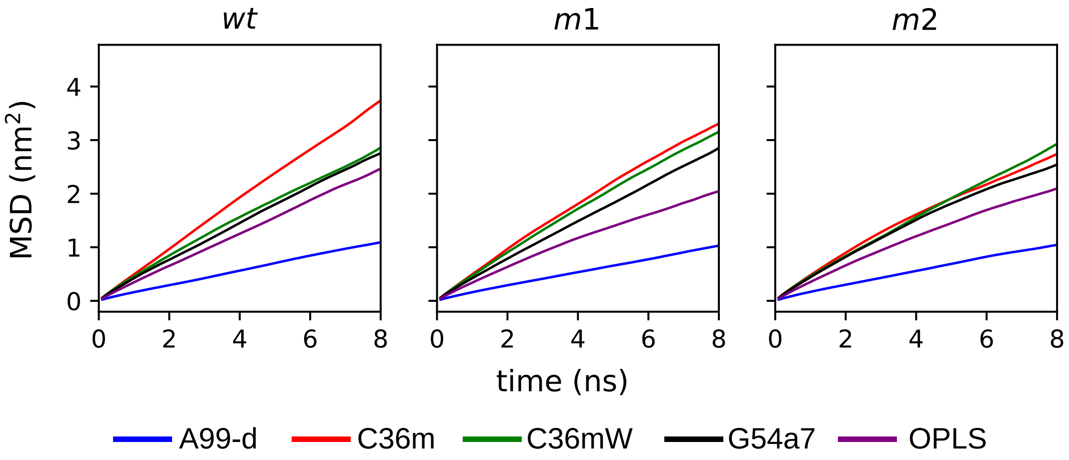


Figure S5: Mean squared displacement (MSD) calculated within 10 ns time windows for $wt$, $m1$, and $m2$. The results are averaged over three independent trajectories. The slope of each curve is measured from 2 ns to 8 ns to obtain the translational diffusion constants.

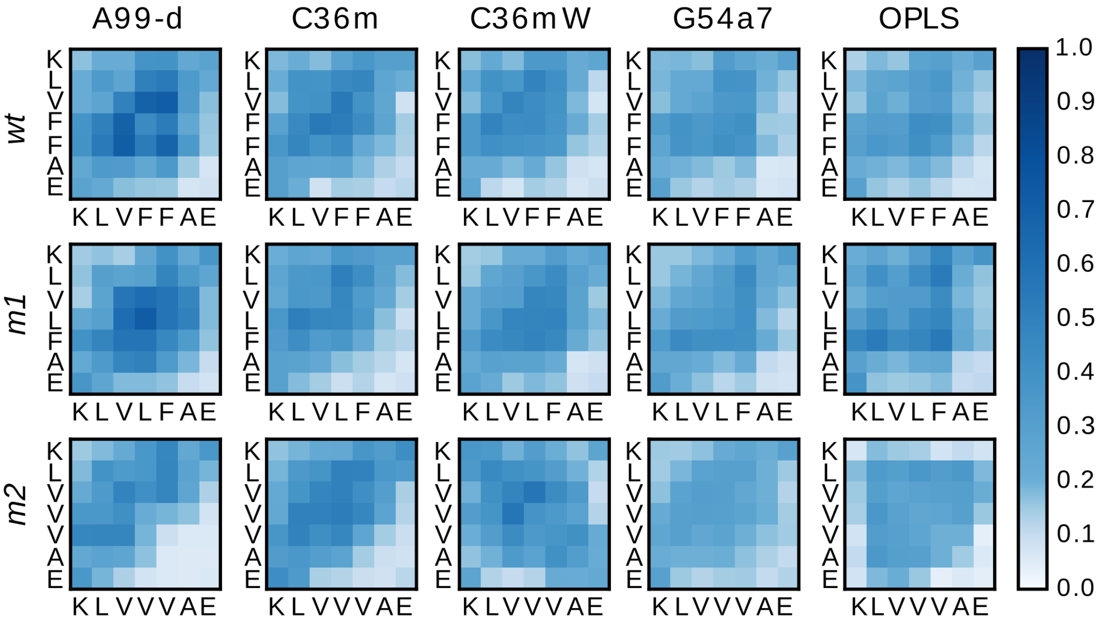


Figure S6: Interpeptide contacts found in the oligomers formed in the simulations of six copies of *wt*, *m*1, and *m*2 using A99-d, C36m, C36mW, G54a7, and OPLS. The color code on the right represents the contact probability between residues.

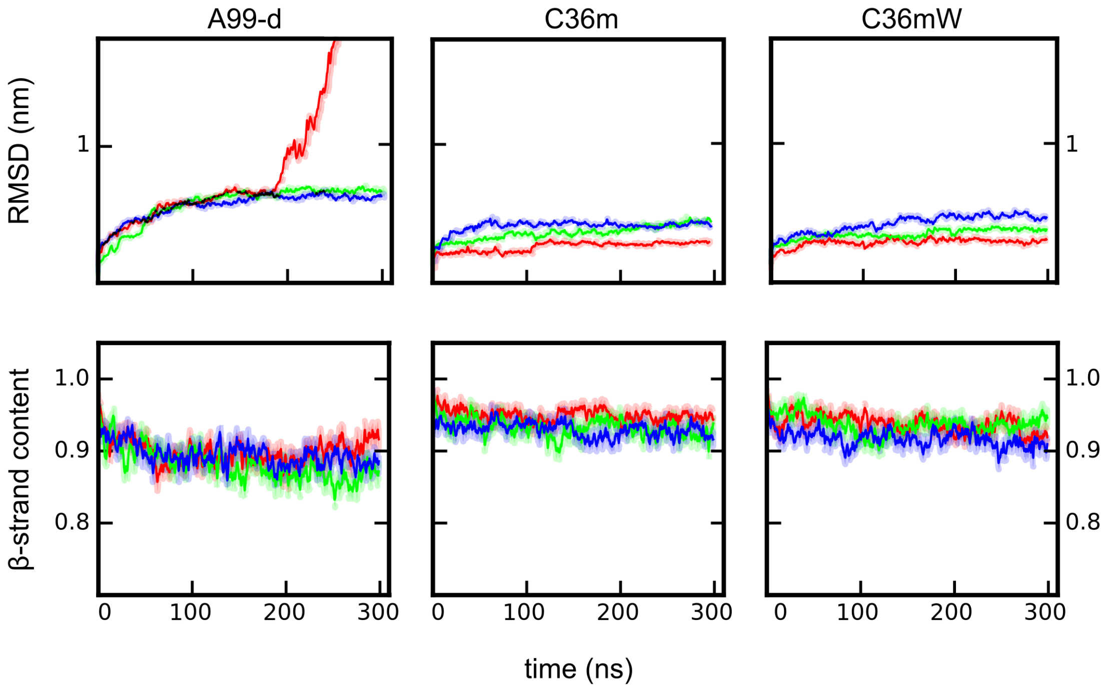


Figure S7: The change in the RMSD (top) and the $\beta$-sheet content (bottom) for the minifibril of *wt* (red), *m*1 (green), and *m*2 (blue) simulated with A99-d, C36m, and C36mW. The averages over three independent simulations per system are shown. The shaded areas indicate the standard error.

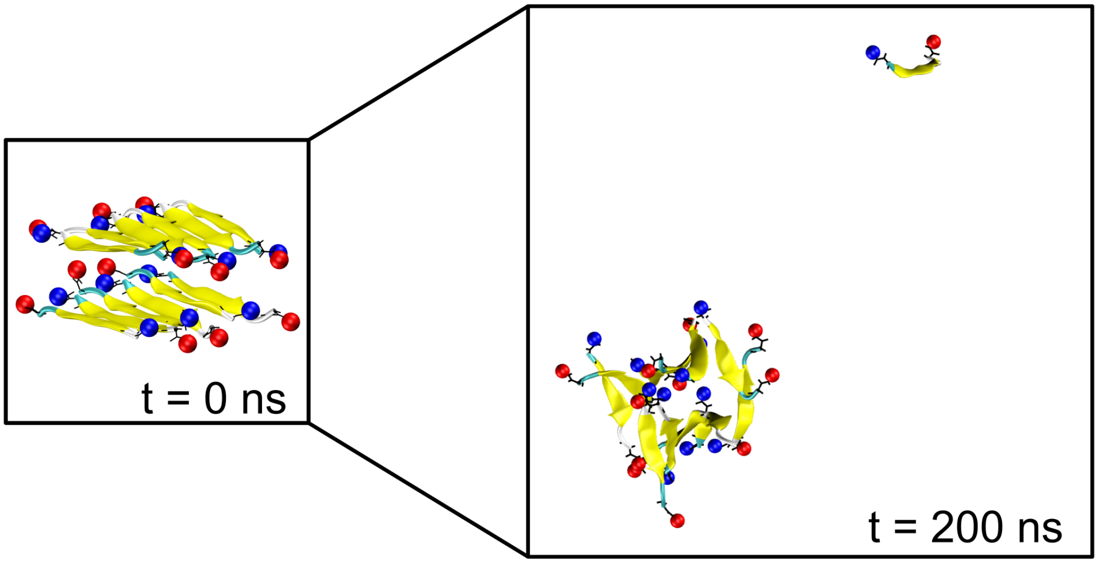


Figure S8: Snapshots from the MD simulation of *wt* using A99-d at $t = 0$ ns and $t = 200$ ns. The representative structure at $t = 200$ ns depicts the detachment of one of the peptides from the fibril.

Table S1: Population (in %) of the first two clusters obtained from structural clustering of the monomer conformations sampled for $wt$, $m1$, and $m2$.

| **Peptide** | **FF** | **Population** | |
|---|---|---|---|
| | | **1$^{st}$ cluster** | **2$^{nd}$ cluster** |
| ***wt*** | **A99-d** | 74% | 12% |
| | **C36m** | 75% | 13% |
| | **C36mW** | 89% | 8% |
| | **G54a7** | 40% | 23% |
| | **OPLS** | 53% | 19% |
| ***m1*** | **A99-d** | 56% | 19% |
| | **C36m** | 64% | 19% |
| | **C36mW** | 72% | 16% |
| | **G54a7** | 49% | 10% |
| | **OPLS** | 43% | 17% |
| ***m2*** | **A99-d** | 66% | 14% |
| | **C36m** | 69% | 15% |
| | **C36mW** | 87% | 8% |
| | **G54a7** | 53% | 16% |
| | **OPLS** | 41% | 19% |

Table S2: Solvent accessible surface area (SASA), number of water molecules, $n_{\mathrm{water}}$, within a cutoff distance of $r_{\mathrm{cut}} = 0.45$ nm with respect to the peptide heavy atoms, and the interfacial water area (IWA), which is the ratio between the SASA and $n_{\mathrm{water}}$, calculated from the simulations of the monomer of $wt$, $m1$, and $m2$, respectively.

| **FF** | **Variant** | **SASA ($nm^2$)** | $\boldsymbol{n}_{\mathrm{water}}$ | **IWA ($nm^2$)** |
|---|---|---|---|---|
| **A99-d** | $wt$ | 13.52 ± 0.03 | 140.66 ± 0.24 | 0.0962 ± 0.0003 |
| | $m1$ | 13.21 ± 0.10 | 137.18 ± 0.83 | 0.0963 ± 0.0009 |
| | $m2$ | 12.68 ± 0.08 | 132.47 ± 0.67 | 0.0957 ± 0.0007 |
| **C36m** | $wt$ | 13.37 ± 0.02 | 138.72 ± 0.10 | 0.0962 ± 0.0003 |
| | $m1$ | 13.19 ± 0.01 | 136.44 ± 0.05 | 0.0967 ± 0.0001 |
| | $m2$ | 12.63 ± 0.04 | 131.26 ± 0.30 | 0.0962 ± 0.0004 |
| **C36mW** | $wt$ | 13.43 ± 0.03 | 140.98 ± 0.31 | 0.0953 ± 0.0003 |
| | $m1$ | 13.26 ± 0.01 | 138.81 ± 0.06 | 0.0955 ± 0.0001 |
| | $m2$ | 12.72 ± 0.02 | 133.76 ± 0.18 | 0.0951 ± 0.0002 |
| **G54a7** | $wt$ | 12.34 ± 0.09 | 128.77 ± 0.77 | 0.0967 ± 0.0009 |
| | $m1$ | 12.14 ± 0.04 | 126.82 ± 0.38 | 0.0957 ± 0.0004 |
| | $m2$ | 11.53 ± 0.02 | 122.51 ± 0.21 | 0.0941 ± 0.0002 |
| **OPLS** | $wt$ | 12.27 ± 0.12 | 127.58 ± 1.08 | 0.0962 ± 0.0013 |
| | $m1$ | 12.14 ± 0.06 | 126.27 ± 0.51 | 0.0962 ± 0.0006 |
| | $m2$ | 11.72 ± 0.02 | 122.29 ± 0.17 | 0.0958 ± 0.0002 |

Table S3: Average electrostatic (Coul) and van der Waals (vdW) interaction energies ($E$), including standard error of the mean, between peptides in oligomers formed by $wt$, $m1$, and $m2$. The sum of both average energy terms is also provided.

| **Peptide** | **FF** | **Non-bonded energy term** | $\boldsymbol{E}$ (kJ/mol) | $\boldsymbol{E}_{\text{Coul}} + \boldsymbol{E}_{\text{vdW}}$ (kJ/mol) |
|---|---|---|---|---|
| ***wt*** | **A99-d** | Coul | $-4.11 \pm 0.29$ | $-9.13$ |
| | | vdW | $-5.02 \pm 0.27$ | |
| | **C36m** | Coul | $-34.30 \pm 3.14$ | $-62.58$ |
| | | vdW | $-28.27 \pm 1.04$ | |
| | **C36mW** | Coul | $-28.33 \pm 3.12$ | $-53.86$ |
| | | vdW | $-25.53 \pm 1.37$ | |
| | **G54a7** | Coul | $-31.00 \pm 1.80$ | $-60.47$ |
| | | vdW | $-29.46 \pm 1.05$ | |
| | **OPLS** | Coul | $-31.04 \pm 2.83$ | $-59.47$ |
| | | vdW | $-28.43 \pm 1.07$ | |
| ***m1*** | **A99-d** | Coul | $-4.55 \pm 0.29$ | $-9.67$ |
| | | vdW | $-5.13 \pm 1.02$ | |
| | **C36m** | Coul | $-34.07 \pm 3.14$ | $-59.26$ |
| | | vdW | $-25.19 \pm 0.98$ | |
| | **C36mW** | coul | $-29.42 \pm 3.12$ | $-54.35$ |
| | | vdW | $-24.92 \pm 1.38$ | |
| | **G54a7** | Coul | $-33.96 \pm 1.80$ | $-61.37$ |
| | | vdW | $-27.41 \pm 0.86$ | |
| | **OPLS** | Coul | $-32.60 \pm 2.83$ | $-57.46$ |
| | | vdW | $-24.86 \pm 1.34$ | |
| ***m2*** | **A99-d** | Coul | $-2.73 \pm 0.88$ | $-5.30$ |
| | | vdW | $-2.57 \pm 0.81$ | |
| | **C36m** | Coul | $-34.25 \pm 4.51$ | $-54.65$ |
| | | vdW | $-20.40 \pm 2.20$ | |
| | **C36mW** | Coul | $-27.13 \pm 2.27$ | $-49.41$ |
| | | vdW | $-22.27 \pm 1.46$ | |
| | **G54a7** | Coul | $-28.21 \pm 2.13$ | $-50.75$ |
| | | vdW | $-22.54 \pm 1.37$ | |
| | **OPLS** | Coul | $-8.05 \pm 0.64$ | $-27.55$ |
| | | vdW | $-19.51 \pm 0.48$ | |
